# AI4Loop: an Artificial Intelligence Framework Reveals Increased 3D Chromatin Interactions and Therapeutic Vulnerabilities across 12,000 Cancer Samples

**DOI:** 10.64898/2026.08.12.744314

**Authors:** Fuying Dao, Benjamin Lebeau, Prachi Amuley, Wei Kit Tan, Xinya Li, Boon Cher Goh, Wee Joo Chng, Chee Keong Kwoh, Hao Lin, Hao Lyu, Melissa Jane Fullwood

## Abstract

Three-dimensional chromatin interactions shape gene regulation, but their large-scale analysis remains limited by the cost and complexity of experimental assays. Here we present AI4Loop, a deep learning framework that infers genome-wide gene-centered chromatin interaction networks directly from RNA-seq data. Across multiple cell types, AI4Loop recovered interaction patterns consistent with clinical samples and orthogonal chromatin conformation datasets. Applied to 12,347 transcriptomes from 32 cancer types, AI4Loop revealed pervasive increases in gene-centered chromatin interactions in tumors, particularly at oncogene-associated loci. These inferred interaction networks outperformed gene expression alone in cancer classification. Integration with more than 50,000 drug-treated transcriptomes identified compounds predicted to reverse cancer-associated interaction gains. Hi-C experiments confirmed that the oxazolidinone antibiotics eperezolid and radezolid reduce breast cancer-gain chromatin interactions. Together, these results identify increased gene-centered chromatin interactions as a pan-cancer feature and provide a scalable strategy for linking 3D genome dysregulation to therapeutic vulnerabilities.

## Introduction

Three-dimensional (3D) genome organization plays a central role in gene regulation by bringing distal genomic loci into spatial proximity and coordinating transcriptional programs across the nucleus (1,2). Despite substantial advances in chromatin conformation capture technologies, systematic characterization of chromatin interaction networks across diverse biological conditions remains challenging due to high cost, technical complexity, and limited scalability (3,4). As a result, much of the 3D regulatory landscape remains inaccessible in large-scale transcriptomic resources generated for human diseases. Here, we focus on gene-centered chromatin interactions (GCIs), defined as chromatin interactions anchored at gene loci, representing spatial coupling between promoter-proximal genomic regions (5) (**Fig. 1A**). Unlike canonical enhancer-promoter interactions that describe specific regulatory links, GCIs provide a higher-level representation of 3D genome organization by capturing coordinated transcriptional relationships among genes within shared nuclear environments (6). Such interactions are thought to organize regulatory hubs that synchronize gene expression programs and contribute to lineage-specific and context-dependent transcriptional control. However, their global organization and functional roles remain poorly understood, particularly across large and heterogeneous biological cohorts (7,8).

**Fig. 1.**
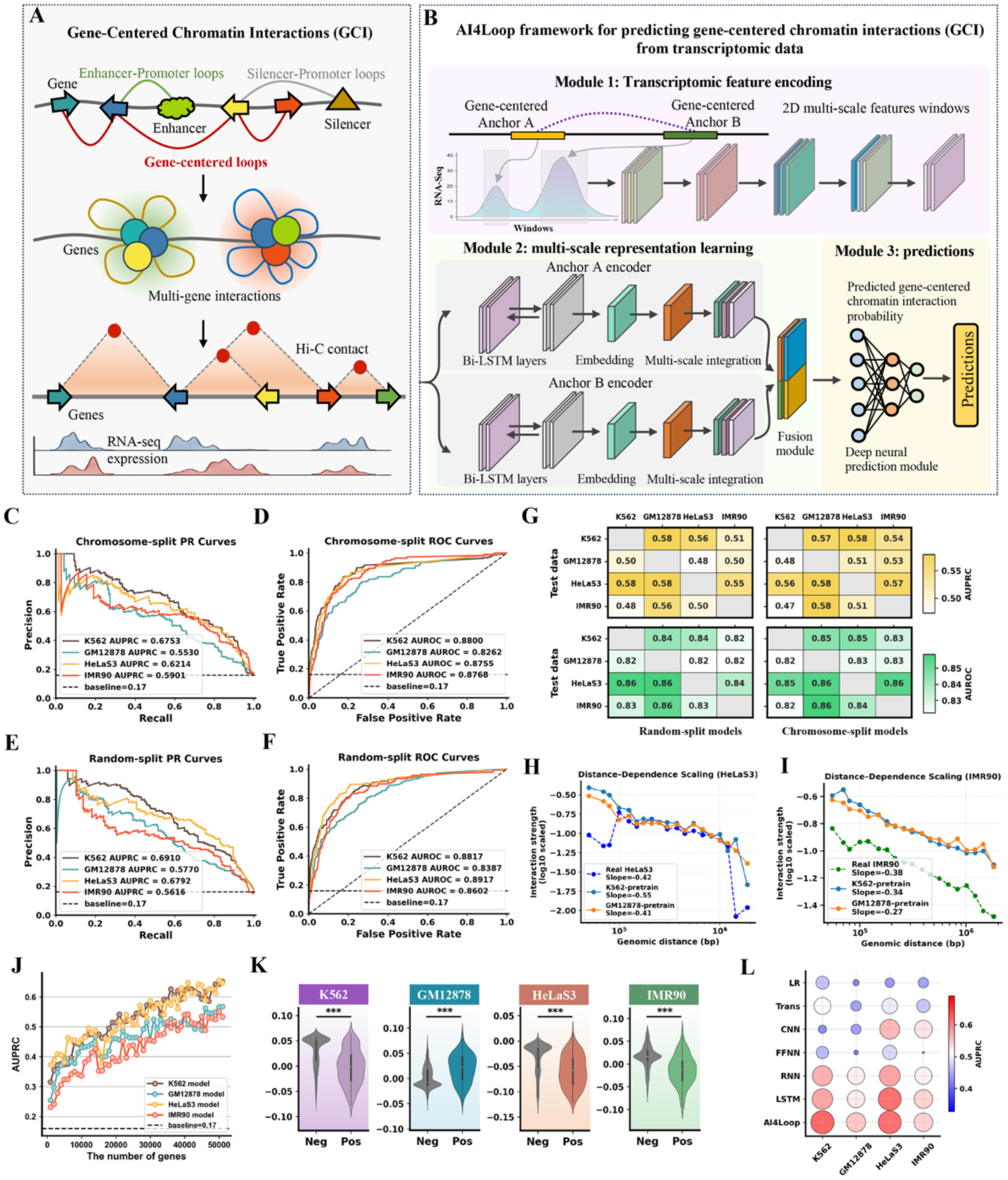
Overview of AI4Loop framework and its performance. (**A**) Gene-centered chromatin interactions (GCIs) represent spatial contacts between promoters of different genes, forming multi-gene regulatory hubs distinct from enhancer-gene loops and silencer-gene loops. (**B**) AI4Loop is a deep learning framework designed to predict GCIs using only RNA-Seq data. Gene expression signals surrounding the anchors of potential interactions are extracted at multiple spatial resolutions and encoded as multi-scale features. These features are input into a bidirectional long short-term memory (Bi-LSTM) model that learns the probability of interaction between gene pairs. (**C-F**) The Precision-Recall (PR) curves and ROC curves for chromosome-split strategy (C and D) and random-split strategy (E and F). (**G**) The heatmap of cross-sample test on random-split and chromosome-split models. The horizontal axis is the model trained on a cell line, and the vertical axis is the test set of other cell lines. (**H-I**) Distance-dependent decay of GCI strength in HeLaS3 and IMR90. Predicted GCIs from both pretrained models recapitulate the polymer scaling behavior observed in real Hi-C data with highly similar decay slopes. (**J**) The model’s AUPRC increases as the number of input genes increases. (**K**) Comparison of the last layers of AI4Loop in positive and negative samples. (L) Comparison of AI4Loop with Transformer (Trans), Convolutional Neural Network (CNN), Feedforward Neural Network (FFNN), Recurrent Neural Network (RNN), LSTM and Logistic Regression (LR) in terms of AUPRC.

Cancer provides a compelling context to investigate this question. Tumor cells exhibit widespread disruption of transcriptional regulation, epigenetic states, and genome architecture (9). yet whether GCIs are systematically altered across cancer types remains unclear. We hypothesized that tumors acquire widespread gains in gene-centered chromatin interactions that reinforce oncogenic transcriptional programs and contribute to malignant phenotypes. Testing this hypothesis, however, is constrained by the limited availability of chromatin conformation data at scale. In contrast, transcriptomic datasets are abundant. Large consortia such as TCGA have generated RNA-seq profiles for thousands of tumors across diverse cancer types, while perturbation resources such as LINCS provide gene expression signatures across a wide range of chemical and genetic perturbations (10). These datasets capture diverse regulatory states, raising the possibility that transcriptomic patterns may encode latent information about the underlying chromatin interaction architecture.

Recent advances in artificial intelligence have enabled the prediction of diverse layers of genome regulation, including gene expression, chromatin accessibility, histone modifications, transcription factor binding and three-dimensional chromatin organization, from DNA sequence and cell-type-specific molecular profiles (11,12). However, many leading models either rely primarily on static sequence information or require matched epigenomic inputs, such as chromatin accessibility and CTCF binding, which are not routinely available in large clinical cohorts and may limit the modelling of sample-specific regulatory variation across biological conditions (13,14). We therefore reasoned that transcriptomic data themselves could provide an alternative entry point for inferring 3D genome organization, as coordinated gene expression patterns may reflect underlying spatial genome structure.

Here, we present AI4Loop, a deep learning framework that infers genome-wide GCI networks directly from RNA-seq data. AI4Loop models multi-scale transcriptional features surrounding candidate gene loci and learns context-dependent relationships between paired regions through a shared bidirectional recurrent architecture, enabling inference of chromatin interaction probabilities from transcriptomic signals alone. Across multiple cell types, AI4Loop recapitulates interaction patterns observed in orthogonal chromatin conformation datasets and clinical samples. Application to more than 12,000 transcriptomes across 32 cancer types revealed a pervasive increase in GCIs in tumors, particularly at oncogene-associated loci. Furthermore, integration with large-scale drug perturbation transcriptomes enabled systematic identification of compounds predicted to reverse cancer-associated interaction gains, with experimental validation using Hi-C. Together, our results demonstrate that transcriptomic programs encode latent information about spatial genome organization and establish a scalable framework for studying chromatin interaction dysregulation and therapeutic vulnerabilities across large biological systems.

## Results

### AI4Loop accurately predicts chromatin interactions using transcriptomic features alone

A central goal of this study was to determine whether GCIs can be inferred directly from RNA-seq data alone. These interactions arise from diverse regulatory configurations and are reflected in coordinated transcriptional activity across interacting loci (**Fig. 1A**). To address this problem, we developed AI4Loop, a deep learning framework for predicting gene-centered chromatin interactions from transcriptomic data (**Fig. 1B**). In AI4Loop, RNA-seq signals surrounding each anchor are encoded using variable-sized sliding windows to capture transcriptional information at multiple spatial scales, and are then processed by shared bidirectional LSTM encoders to model dependencies within each locus while generating comparable representations for paired anchors. These anchor-level features are subsequently integrated through a nonlinear prediction module to estimate chromatin interaction probabilities. Through this design, AI4Loop captures structured and context-dependent transcriptional patterns associated with chromatin looping, providing a transcriptome-based framework for inferring gene-centered chromatin interactions.

To evaluate the performance of AI4Loop across multiple cell lines, we employed two training strategies: random-split and chromosome-split. The random-split approach was used to assess overall performance, while chromosome-split strategy was designed to eliminate chromosomal dependencies between the training and test sets, thereby mitigating performance inflation (15,16). Random-split achieved AUPRCs > 0.5871 and AUROCs > 0.8199, and chromosome-split maintained AUPRCs > 0.5363 and AUROCs > 0.8467, well beyond the baseline (AUPRC = 0.17, AUROC = 0.5) (**Fig. 1C-F**). These results indicate that AI4Loop exhibits robust predictive performance and generalizes effectively across chromosomes.

To further examine model generalizability, we evaluated cross-cell predictions by training on one cell line and testing on others. AI4Loop maintained strong performance with AUPRC > 0.47 and AUROC > 0.82 (**Fig. 1G**, **Fig. S1**). Extending this analysis, models trained on K562 or GM12878 generalized well to HeLaS3 and IMR90, achieving comparable AUPRC values across cell types (**Fig. S2A-F**). The predicted scores further exhibited distance-dependent decay patterns closely matching Hi-C measurements in HeLaS3 and IMR90 (**Fig. 1H-I**), indicating that AI4Loop captures biologically coherent loop signals (**Fig. S2G**). To rule out inflation from shared loops, we evaluated strictly cell-type–specific interactions (**Fig. S3A-D**). Even under this stringent setting, AI4Loop achieved AUPRCs of 0.53-0.73 and AUROCs of 0.84-0.89 (**Fig. S3F-I**), demonstrating strong transferability and robust generalization to loops uniquely present in the target cell type.

To investigate how input feature design influences AI4Loop performance, we examined the impact of input gene quantity and gene set composition. Model performance increased with the number of input genes, reaching optimal accuracy when genome-wide RNA-seq profiles (>60,000 genes) were used (AUPRC > 0.55; **Fig. 1J**), whereas restricting inputs to small gene sets markedly reduced performance (e.g., ∼0.23 AUPRC using 1,000 genes). Additional analyses further showed that gene features derived from long genes, cancer-associated genes, highly expressed genes, and enhancer-linked genes were generally more informative for chromatin interaction prediction than random or less functionally relevant gene sets (**Fig. S4A-D**). Additional analyses confirmed that AI4Loop benefits from multi-scale, anchor-centered transcriptional representations, with mean-based aggregation yielding the most robust performance, whereas genomic distance alone contributed little predictive power and did not materially improve RNA-based predictions (**Fig. S4E-J**). These results highlight that AI4Loop derives its predictive power primarily from biologically informative, multi-scale transcriptional signals rather than simple distance effects.

To interpret the features learned by AI4Loop, we examined the output features from the last layer of AI4Loop, which showed significant differences between positive and negative samples across all four cell lines (t-test, *p* < 0.001; **Fig. 1K**), indicating that the model learns discriminative transcriptional representations associated with chromatin interactions. Consistently, analyses across individual sliding-window settings showed that 6-7 kb windows were the most informative, while the overall performance trends remained highly consistent across different window sizes (**Fig. S5A-D**). Notably, *NPM1*, a well-established leukemia-associated gene, was identified as a prominent positive feature in K562, supporting the relevance of AI4Loop-derived signals to leukemia biology. Likewise, *HBE1*, one of the key hemoglobin genes, also showed strong contribution in K562, consistent with the erythroid character of this cell line (**Fig. S5E-F**). These results indicate that AI4Loop is not merely a predictive black box, but instead learns biologically meaningful and spatially structured transcriptional features that underpin chromatin interactions. Finally, benchmarking against a range of classical machine learning and deep learning models showed that AI4Loop consistently outperformed all alternatives across all four cell lines (**Fig. 1L**, **Fig. S6**), highlighting the advantage of its architecture in capturing informative patterns from transcriptomic data associated with chromatin interactions.

### AI4Loop generalizes to clinical samples and orthogonal chromatin interaction platforms

We next asked whether AI4Loop could generalize beyond curated cell-line datasets to clinical transcriptomes and independent chromatin conformation platforms. To systematically evaluate AI4Loop’s performance in clinical settings, we tested it on RNA-Seq and Hi-C data from six Chronic Lymphocytic Leukemia (CLL) patient samples (17). Using the pre-trained model from the near-normal cell line GM12878, AI4Loop achieved prediction accuracies ranging from 61% to 67% across samples, which substantially exceeded the random baseline of 50% when compared with matched Hi-C loops (**Fig. 2A** left). Notably, the model trained on the cancer cell line K562 exhibited even higher accuracy, consistently exceeding 88% (**Fig. 2A** right), demonstrating superior predictive performance in cancer-derived transcriptomes.

**Fig. 2.**
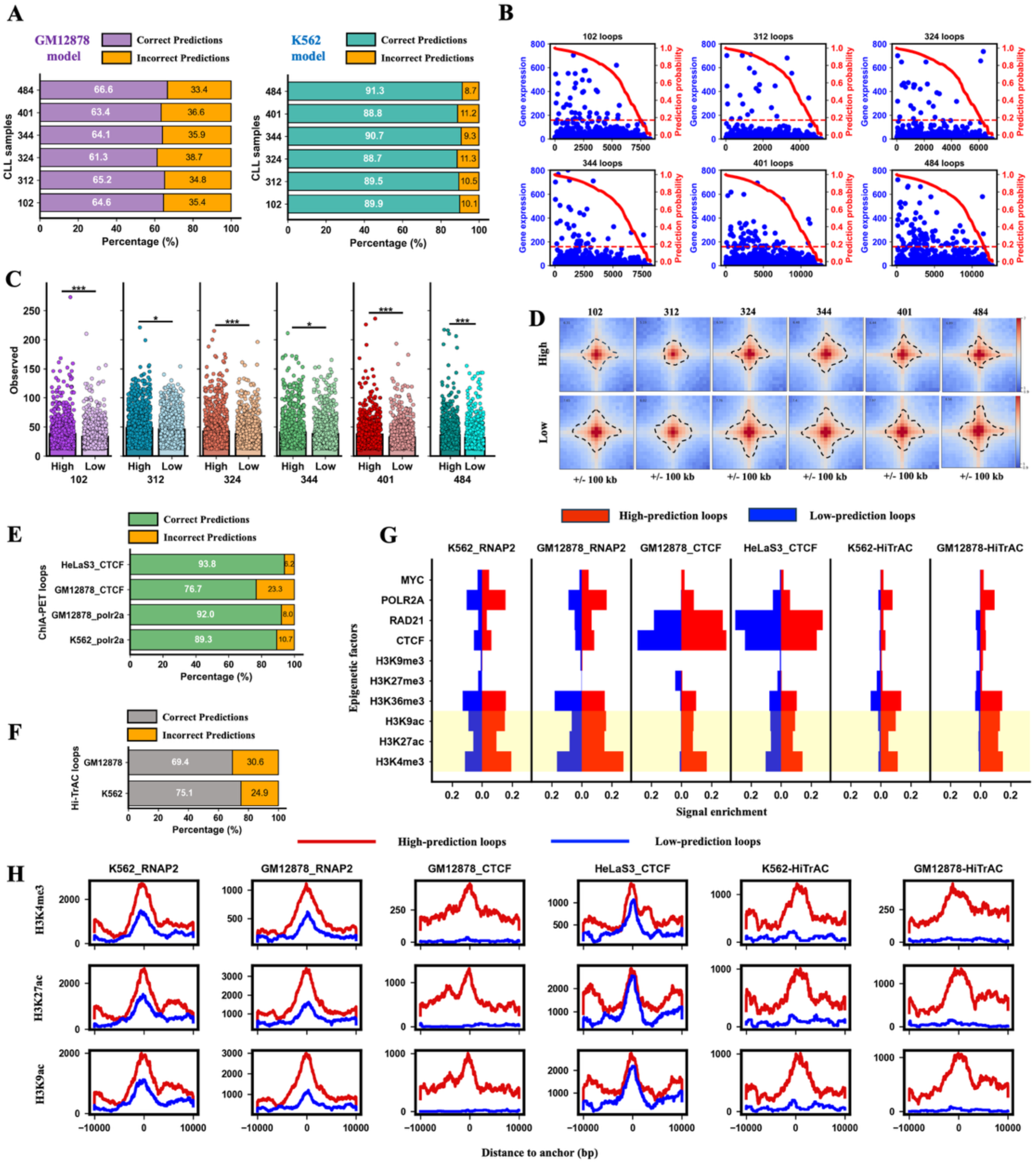
AI4Loop application on clinical data and other data of higher-resolution chromatin conformation capture technologies. (**A**) GM12878 and K562 AI4Loop models respectively were used to predict the Hi-C loops of six Chronic Lymphocytic Leukemia (CLL) samples. (**B**) Displays the predicted probabilities and gene expressions levels of CLL loops. Chromatin loops with high predicted probabilities have more significant gene expression signals. (**C**) Comparing the real Hi-C observed values (the observed value indicates the strength of the real Hi-C loop) with high and low prediction probabilities. It shows that the high prediction probability matches the strong loops, indicating that our AI4Loop model is reliable. (**D**) Aggregate peak analysis (APA) shows high-predicted probabilities loops have a strong signal concentration, with peaks being more centralized. High: 1,000 high-predicted probabilities loops, Low: 1,000 low-predicted probabilities loops. (**E**) Model to predict ChIA-PET loop data of K562-RNAP2, GM12878-RNAP2, GM12878-CTCF and HeLaS3-CTCF. RNAP2 means RNA Polymerase II. (**F**) Model to predict Hi-TrAC loop data. (**G**) The enrichment of epigenetic modification signals in loops with high and low prediction probabilities. (**H**) The distribution of histone modification signals at high and low loop anchors.

Further analysis revealed that gene pairs with higher prediction probabilities were associated with stronger expression signals (**Fig. 2B**), and their expression levels were significantly higher than those of low-confidence predictions (**Fig. 2C**), aligning well with real-world data. Aggregate Peak Analysis (APA) (18) of Hi-C data further confirmed these predictions: high-confidence interactions exhibited narrower, more concentrated interaction peaks than low-confidence ones (**Fig. 2D**). Consistently, when stratified by genomic distance, correct prediction ratios decreased gradually with increasing loop distance but remained stably above random expectation across all distance ranges (**Fig.S7A-B**), and high-confidence predictions showed strong concordance with focal interaction patterns in representative Hi-C comparison regions (**Fig. S7C**).

To confirm that AI4Loop predictions are not driven by Hi-C induced biases, we further tested AI4Loop on orthogonal chromatin interaction datasets, including ChIA-PET and Hi-TrAC (19,20). Across these platforms, AI4Loop maintained high accuracy, consistently exceeding 70%. Notably, the HeLaS3-trained model achieved an accuracy of 93.8%, while the GM12878 model correctly predicted 92% of POLR2A-associated interactions (**Fig. 2E-F**). High-confidence interactions were enriched for active chromatin features, including strong transcriptional signals and the presence of active enhancers and promoters (**Fig. 2G-H, Table S1**), indicating that AI4Loop effectively captures transcriptionally relevant chromatin architecture. Conversely, repressive histone marks such as H3K9me3 and H3K27me3 were depleted in these high-confidence predictions (**Fig. 2G**), suggesting that most predicted GCIs are associated with gene activation.

Collectively, these results demonstrate that AI4Loop not only performs robustly across multiple 3D genomics platforms, but also generalizes effectively to clinical RNA-Seq samples from previously unseen patients. The strong concordance between AI4Loop predictions and experimentally observed chromatin features highlights its potential as a scalable, sequencing-efficient tool for decoding chromatin interactions in real-world disease contexts, including personalized transcriptomes and drug response profiling in cancer.

### AI4Loop reveals leukemia-associated chromatin interaction programs in Acute Myeloid Leukemia

We next investigated whether AI4Loop could uncover disease-associated chromatin interaction programs in acute myeloid leukemia (AML). Using a genome-wide K562-trained model, we first focused on the highest-confidence predicted GCIs among 3,590,596 candidate intrachromosomal gene pairs (**Fig. S8A**). Compared with low-confidence pairs, the top 1,000 predicted GCIs were strongly enriched for super-enhancers, active histone marks (H3K4me3, H3K27ac, H3K9ac, and H3K4me1), and chromatin architectural regulators including CTCF, RAD21, POLR2A, and YY1 (**Fig. 3A**, **Fig. S8D**). A subset also overlapped super-silencers and H3K27me3-marked regions or displayed bivalent signatures, consistent with poised regulatory states (**Fig. 3B**, **Fig. S8B**, **Table S2**). When applied to AML and healthy samples, these high-confidence GCIs showed marked gains in AML, particularly at oncogene-associated loci (**Fig. 3C-D**). From these comparisons, we identified 71 AML-specific and 27 healthy-specific GCIs (**Fig. 3E**, **Table S3**), and hierarchical clustering based on these interactions sharply separated AML from healthy samples (ARI = 0.8988; NMI = 0.8531; **Fig. 3F**).

**Fig. 3.**
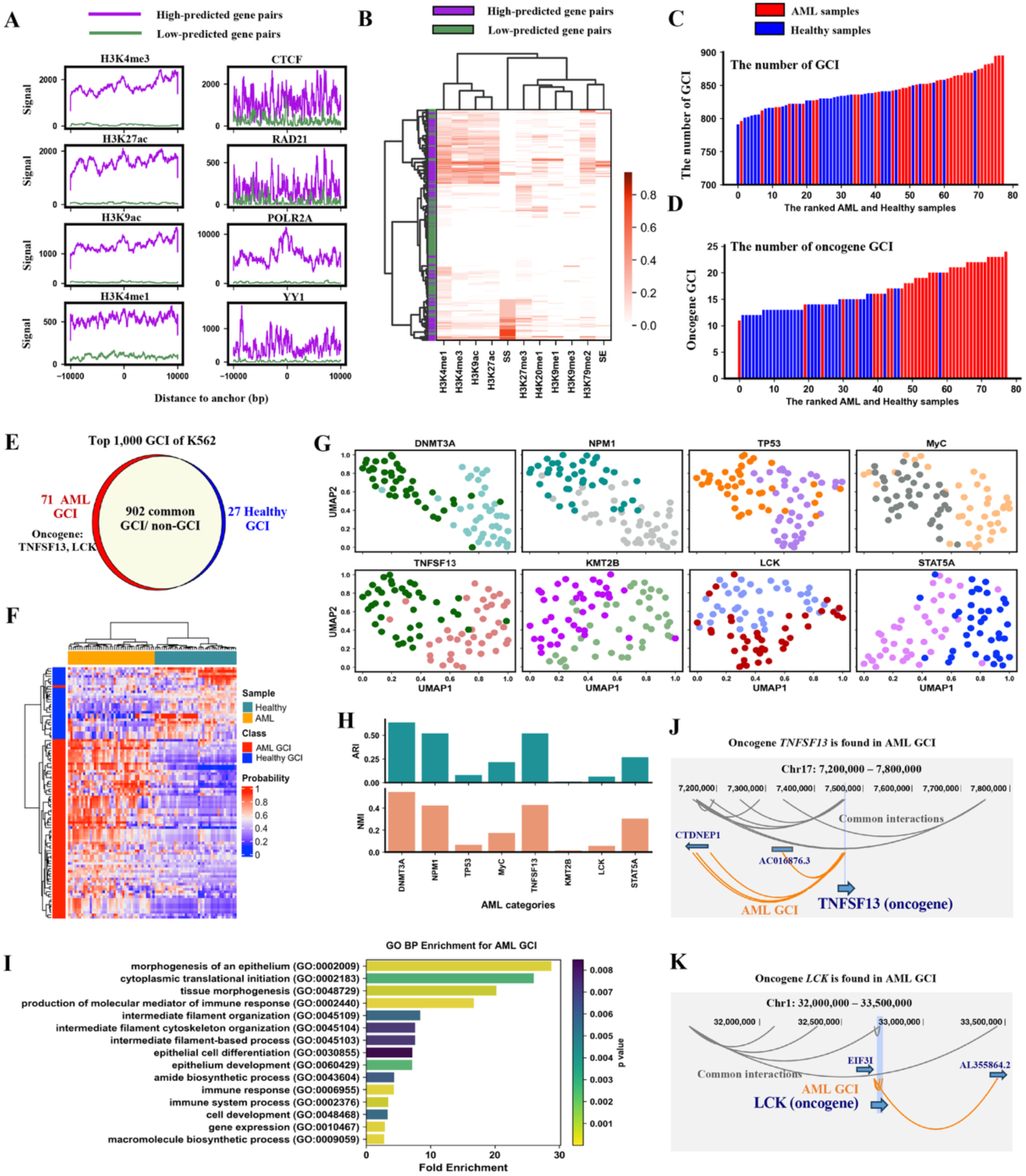
AI4Loop revealed the GCI differences between different AML samples and subtypes. (**A**) The distribution of epigenetic modification signals at K562 gene pairs with high and low prediction probabilities. (**B**) Hierarchical clustering high- and low- prediction gene pairs base on the enrichment of epigenetic modification signals. (**C**) The number of GCIs in AML and healthy samples. (**D**) The number of oncogene-GCIs in AML and healthy samples. (**E**) Venn diagram showing the number of AML GCI, Healthy GCI and Common GCI among the top 1,000 GCI of K562. (**F**) Hierarchical clustering in AML (orange) and healthy bone marrow samples (cyan) based on prediction probabilities of 71 AML GCIs (red) and 27 Healthy GCIs (blue). (**G**) Clustering of subtypes of AML by UMAP based on predicted top 1,000 GCIs of K562. For example, we have high-*DNMT3A* and low-*DNMT3A* AML samples according to the Fragments Per Kilobase of transcript per Million mapped reads (FPKM) of DNMT3A. Here we show ten subtypes of AML. (**H**) Evaluation of AML subtype clustering performance using Adjusted Rand Index (ARI) and Normalized Mutual Information (NMI). ARI ranges from -1 (no agreement) to 1 (perfect agreement). NMI ranges from 0 (no mutual information) to 1 (perfect mutual information). (**I**) Gene Ontology (GO) functional analysis of AML GCIs. (**J** and **K**) Visualization of AML oncogene GCIs, including *TNFSF13* and *LCK* genes.

AML-associated GCI profiles also captured subtype-related structure across patient samples. Using RNA-seq data from 151 patients in the TCGA-LAML cohort, we grouped samples according to the expression of established AML-related genes, including *DNMT3A*, *NPM1*, *TP53*, and *MYC*, as well as additional interaction-associated genes identified by AI4Loop, including *TNFSF13*, *KMT2B*, *LCK*, and *STAT5A*. UMAP projections based on AI4Loop-predicted chromatin interactions revealed particularly strong separation in *DNMT3A*-, *NPM1*-, and *TNFSF13*-associated groups, whereas TP53- and MYC-associated groups showed weaker stratification (**Fig. 3G-H**). Consistent with these results, genes involved in AML-specific GCIs were significantly enriched in biological processes central to leukemia pathogenesis, including cell differentiation, immune response, cytoskeletal organization, biosynthesis, and gene regulation, whereas healthy-specific GCIs were enriched in processes related to cellular homeostasis (21) (**Fig. 3I**, **Fig. S8C**).

Finally, AML-specific GCIs highlighted candidate cooperative oncogenic relationships that were not apparent from single-gene analyses alone. Two illustrative examples were the AML-specific GCIs linking *TNFSF13* with *CTDNEP1* and *LCK* with *EIF3I* (**Fig. 3J-K**). *TNFSF13* is associated with pro-proliferative *NF-κB* and *PI3K–AKT* signaling, whereas *CTDNEP1* contributes to nuclear envelope stability, raising the possibility that their coordinated chromatin coupling links growth signaling with stress tolerance. Similarly, the *LCK–EIF3I* interaction connects mTOR-related signaling with oncogenic translational control, suggesting coordinated promotion of aberrant protein synthesis. Together, these findings indicate that AML is characterized by increased oncogene-associated chromatin interactions and that AML-specific GCI programs can reveal candidate regulatory dependencies not readily apparent from transcript-level analyses alone.

### AI4Loop enables scalable pan-cancer mapping of chromatin interaction states from RNA-seq

We next asked whether AI4Loop could scale from benchmark datasets to large clinical transcriptome cohorts and reveal pan-cancer chromatin interaction patterns. To address this question, we applied AI4Loop to a pan-cancer transcriptome compendium comprising 12,347 TCGA RNA-seq samples, including 10,042 tumors and 2,305 normal tissues across 32 cancer types (**Fig. 4A-C**). From 5,338 consistently expressed genes, we constructed 49,255 intrachromosomal gene pairs and predicted their probabilities using pre-trained GM12878 model with threshold ≥ 0.5 (**Fig. S9A**). Among these, 8,398 pairs were consistently non-interacting across all samples (prediction score < 0.5, 17.05%), while the remaining 40,857 pairs exhibited sample-specific interaction patterns (82.95%) and were further organized into distinct categories based on their prevalence across samples, including GCI-all (shared across all samples, n = 855, 1.73%), GCI-cancer (enriched in cancer, n = 101, 0.21%), GCI-healthy (enriched in healthy tissues, n = 17, 0.03%), GCI-cancer-one (restricted to a single cancer type, n = 1,835, 3.73%), and GCI-several (present across several but not all cancer types, n = 38,049, 77.25%) (**Fig. 4D-E**). Notably, universally detected interactions (GCI-all) included canonical housekeeping gene pairs such as *GAPDH* and *B2M* (22), supporting the biological plausibility of the inferred interaction map.

**Fig. 4.**
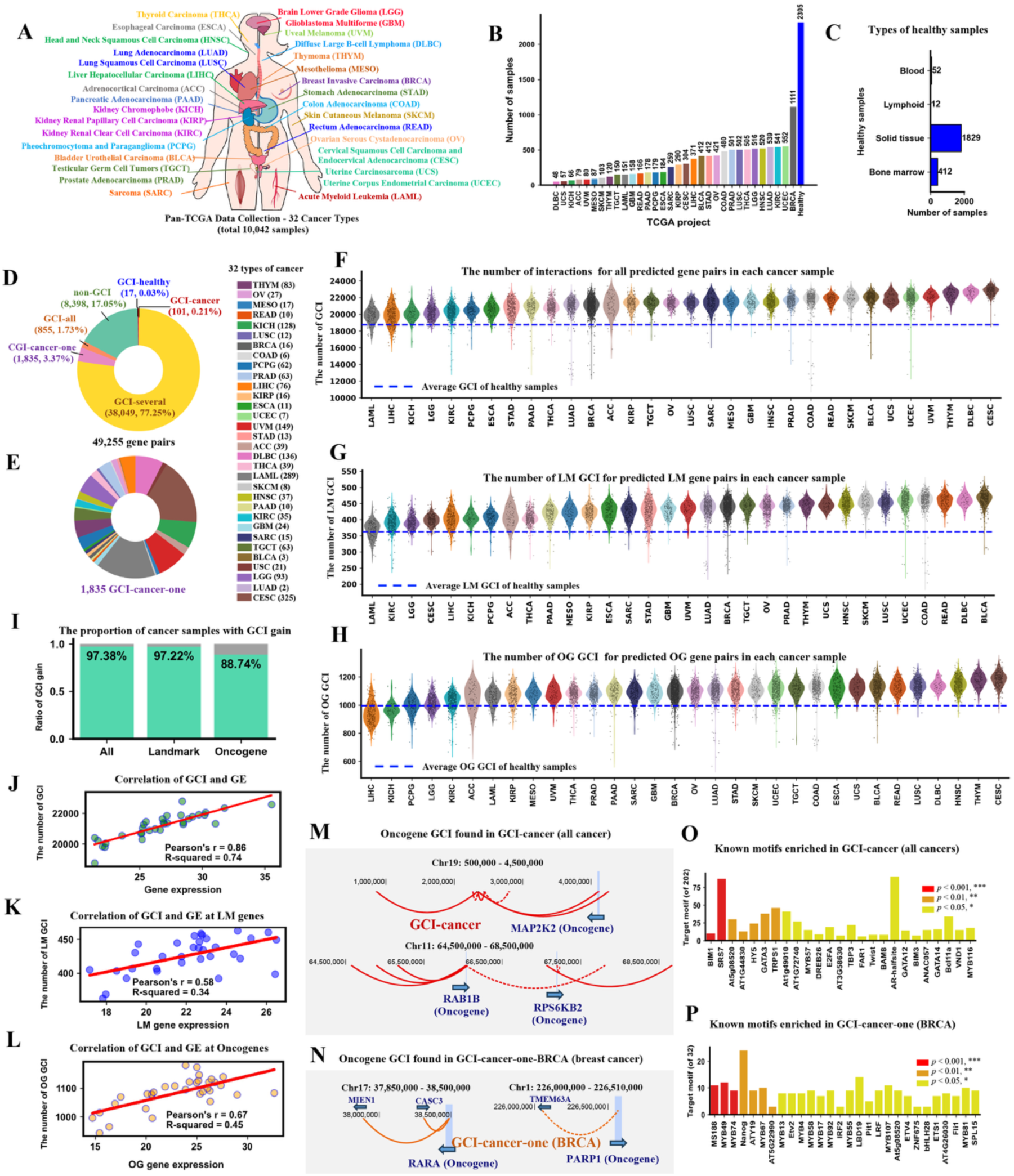
AI4loop reveals that cancers have increased GCIs as compared with healthy samples. (**A**) Pan-cancer RNA-Seq data was collected from The Cancer Genome Atlas database (TCGA). (**B**) The number of each type of cancer and healthy samples. (**C**) The types and numbers of healthy samples. (**D**) 49,255 gene pairs were divided into non-GCI (17.05%) and GCI (82.95%) according to the predicted probability of AI4Loop, among which GCIs were further divided into GCI-all, GCI-cancer, GCI-healthy, and GCI-cancer-one. (**E**) GCI-cancer-one is the GCI unique to each type of cancer. (**F-H**) The number of GCIs for all predicted 49,255 gene pairs, 1,290 landmark (LM) gene pairs, and 4,090 oncogene (OG) pairs. (**I**) Proportion of cancer samples with increased GCIs among the predicted gene pairs. (**J-L**) Correlation between GCI counts and average gene expression levels. (J) All genes, (K) oncogenes, and (L) landmark genes, with Pearson’s r = 0.86, 0.67, and 0.58 and R² = 0.74, 0.45, and 0.34, respectively. (**M**) GCI-cancer at oncogene *MAP2K2*, *RAB1B*, and *RPS6KB2*. (**N**) GCI-cancer-one-BRCA (breast cancer) at oncogene *PARP1* and *RARA.* (**O**) Enriched motifs of GCI-cancer. (**P**) Enriched motifs of GCI-cancer-one-BRCA.

We next focused on the global interaction levels differed between tumors and healthy tissues. Across all predicted 49,255 gene pairs, 97.38% of tumors exhibited increased numbers of GCI relative to healthy controls (**Fig. 4F**). A similar pattern was observed when the analysis was restricted to 1,290 interactions derived from CLUE landmark genes, for which 97.22% of tumors showed elevated interaction frequencies (**Fig. 4G**). When focusing on 4,090 oncogene-associated gene pairs, 88.74% of tumors likewise displayed increased interaction levels relative to healthy tissues (**Fig. 4H**). Across all three interaction sets, GCI abundance was positively correlated with transcript abundance, including global gene expression, landmark gene expression, and oncogene expression, respectively (**Fig. 4J-L**, **Fig. S9B-D**). These results indicate that widespread gains in GCI are a pervasive feature of cancer and are closely linked to elevated transcriptional activity.

We then examined the biological specificity of these cancer-associated interaction gains at oncogene-associated loci. The inferred interaction landscape included both pan-cancer recurrent programs, such as those involving *MAP2K2*, *RAB1B*, and *RPS6KB2*, and cancer-type-restricted oncogenic networks, including *BRCA1*–*KRT9/13/17* interactions in cervical cancer and *TNFSF13*/*PDGFRB* interactions in leukemia (**Fig. 4M-N**, **Fig. S9E-F**). These patterns indicate that GCI gains are not randomly distributed across the genome but preferentially involve loci with established oncogenic relevance. The BRCA1-KRT interaction network is particularly notable, as both genes have been independently associated with basal-like phenotypes and poor prognosis (23,24), suggesting that their coordinated chromatin coupling may contribute to cancer-specific regulatory programs.

To further explore potential regulatory mechanisms underlying these interaction gains, we performed motif enrichment analysis on cancer-enriched GCI sets and identified transcription factor families including GATA, MYB, and KLF as candidate regulators (**Fig. 4O-P**, **Fig. S9G-H**). Because these factors are known to regulate proliferation, invasion, and lineage-specific transcription, their enrichment suggests that tumor-associated GCI gains may be actively maintained by pro-tumorigenic regulatory circuits (25-27). Together, these findings indicate that oncogene-associated GCI gains define structured regulatory programs in cancer and may represent candidate epigenetic vulnerabilities. To facilitate further investigation of these structure-function relationships and the discovery of additional oncogenic synergies, we provide the pan-cancer chromatin interaction atlas generated in this study as a community resource through the Nanyang Technological University data repository (https://doi.org/10.21979/N9/ORBU74).

### AI4Loop-derived chromatin interactions improve cancer-state stratification beyond RNA expression

We next asked whether AI4Loop-derived GCI profiles provide a more informative representation of cancer-state variation than RNA expression-based features. To address this, we compared clustering based on AI4Loop-predicted GCI with clustering based on gene expression or gene co-expression in three complementary settings: cancer-type-specific interactions (GCI-cancer-one), pan-cancer versus healthy interactions (GCI-cancer and GCI-healthy), and landmark gene-restricted interactions. In all three settings, GCI-based clustering produced markedly clearer sample separation than either expression- or co-expression-based clustering (**Fig. 5**). Using cancer-type-specific interactions, AI4Loop-derived GCI profiles separated healthy tissues and diverse cancer types more clearly than the corresponding gene expression or co-expression features (**Fig. 5A-C**). Likewise, cancer-enriched and healthy-enriched GCI sets robustly separated pan-cancer and healthy samples, whereas expression-derived features showed substantially weaker resolution (**Fig. 5D-F**).

**Fig. 5.**
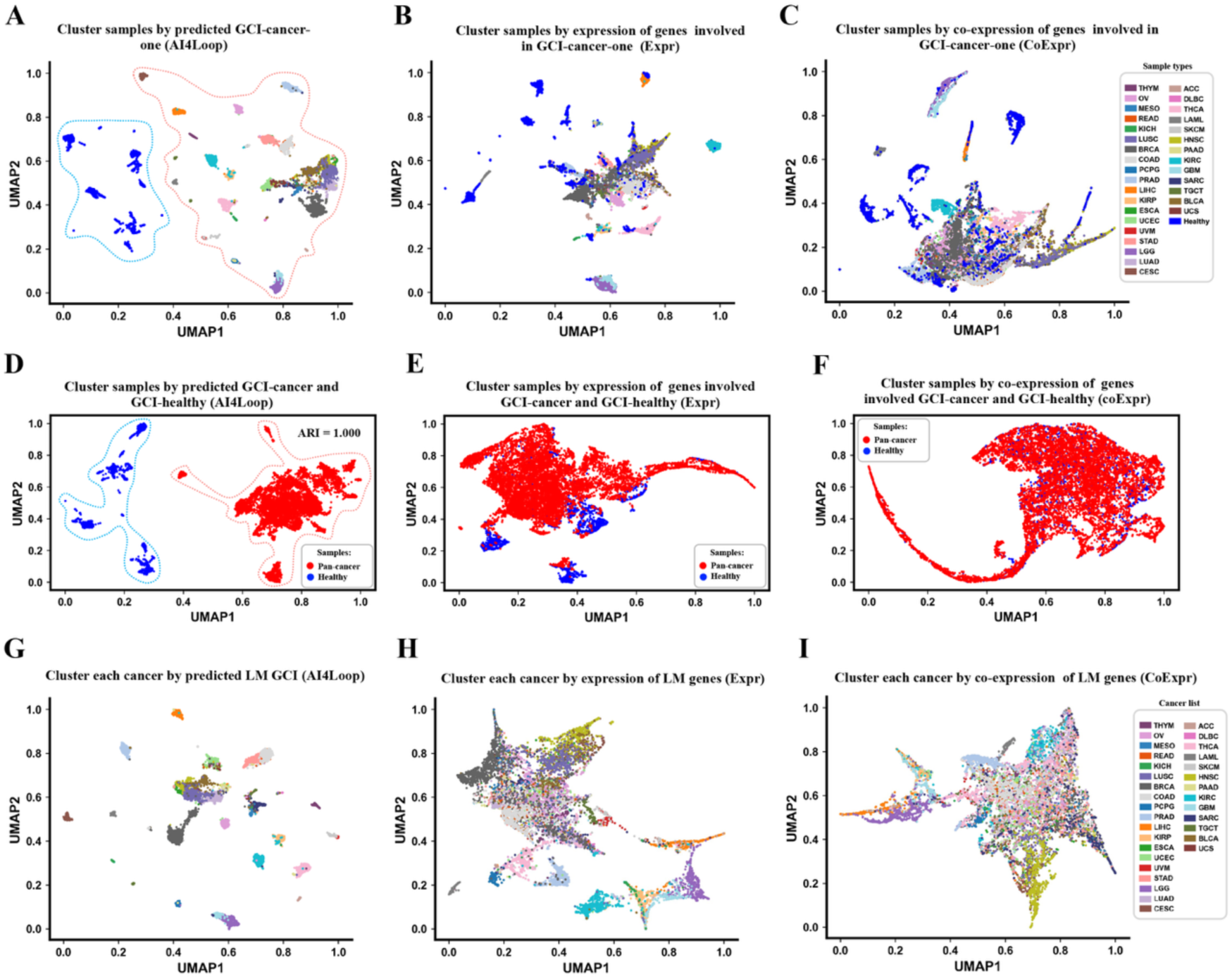
Gene-centered chromatin interactions provide a superior representation for distinguishing cancer states. (**A-C**) UMAP projections of samples based on GCI-cancer-one features. (A) AI4Loop-predicted GCI probabilities. (B) Expression of genes involved in GCI-cancer-one. (C) Co-expression of genes involved in GCI-cancer-one. (**D-F**) UMAP projections of pan-cancer and healthy samples based on GCI-cancer and GCI-healthy features. (D) AI4Loop-predicted GCI probabilities. (E) Expression of genes involved in GCI-cancer and GCI-healthy. (F) Co-expression of genes involved in GCI-cancer and GCI-healthy. (**G-I**) UMAP projections of samples based on landmark gene GCI features. (G) AI4Loop-predicted landmark GCI probabilities. (H) Landmark gene expression. (I) Landmark gene co-expression.

This advantage remained evident even under a reduced feature space restricted to landmark genes, where AI4Loop-predicted landmark GCI still retained strong separation among cancer types, while landmark gene expression and co-expression remained diffuse and poorly resolved (**Fig. 5G-I**). Quantitative evaluation further confirmed the superiority of GCI-based representations across all three settings (**Fig. S10**), with consistently higher clustering consistency and separation metrics than expression- or co-expression-based features. These results show that GCI defines a more structured and biologically meaningful representation of transcriptomes, enabling improved discrimination of healthy and cancer samples as well as finer stratification among cancer types.

### AI4Loop-based drug perturbation profiling identifies compounds targeting oncogenic chromatin interactions

To explore whether transcriptome-inferred chromatin interaction states can be leveraged to identify therapeutically relevant perturbations, we applied AI4Loop to 57,353 drug-induced transcriptomic profiles across 42 cell lines from the CLUE database (**Fig. 6A**). Using the GM12878-pretrained model, we constructed a large-scale perturbation atlas of GCI. Given our observation that cancers exhibit widespread gains in GCI, we hypothesized that compounds capable of reducing these interaction programs may counteract oncogenic transcriptional states. Across cell lines, most perturbational conditions were associated with increased predicted GCI, whereas interaction-reducing responses were relatively depleted in cancer cell lines but more frequently observed in non-malignant contexts (**Fig. 6B-C**), suggesting that reinforcement of chromatin interaction programs may represent a common transcriptional response that is further amplified in cancer.

**Fig. 6.**
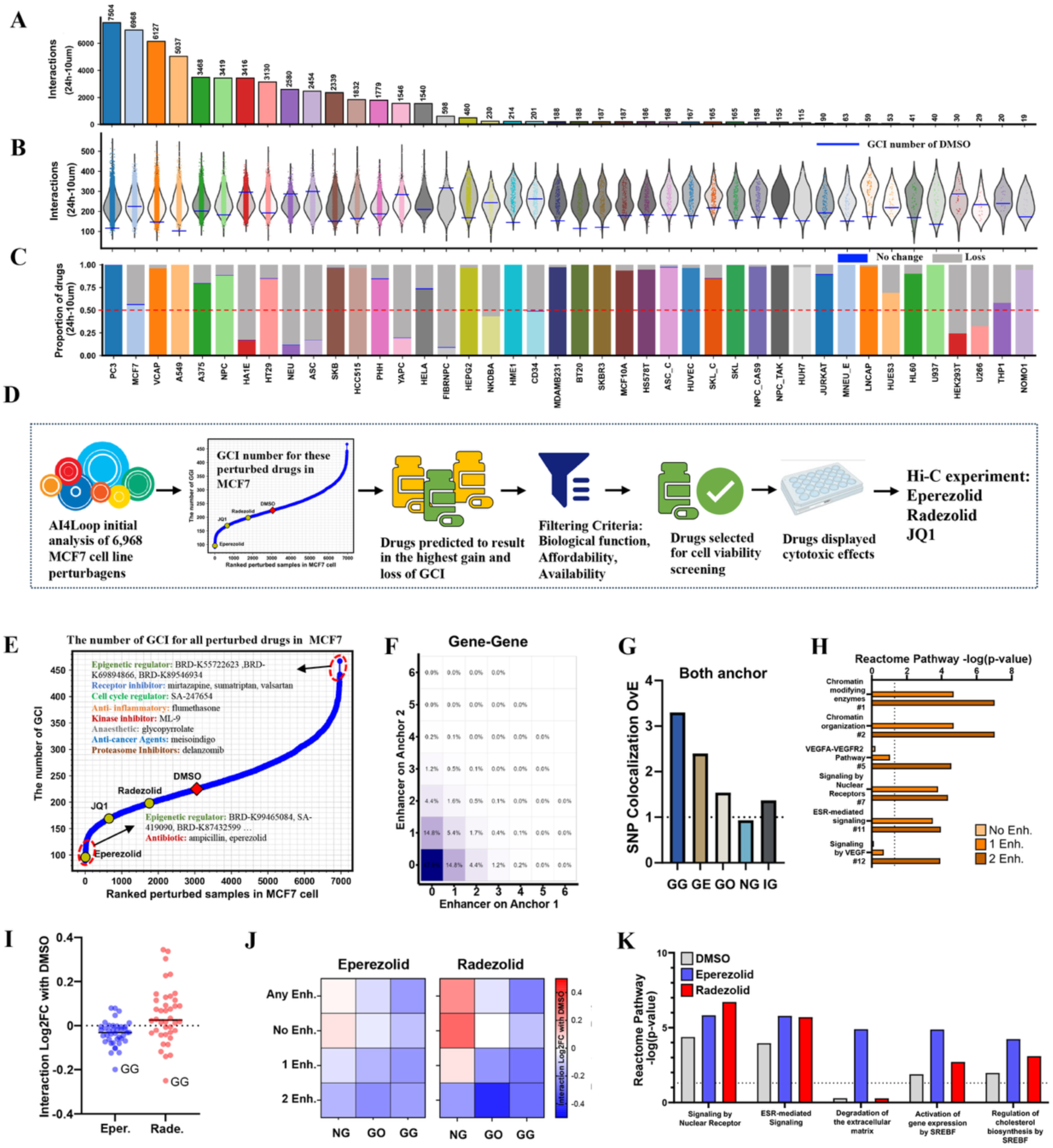
Identifies compounds that modulate gene-centered chromatin interactions (GCIs) using drug-perturbation data from the CLUE database. (**A**) Number of perturbagens tested across 42 cell lines in the CLUE dataset. (**B**) Summary of global GCI gain or loss across cell lines after drug treatment. (**C**) Proportions of GCI gain, loss, and no-change events across all conditions. (**D**) Overview of the AI4Loop-based drug screening workflow. AI4Loop was applied to predict GCI in drug-treated MCF7 samples; compounds inducing the strongest GCI loss were prioritized, filtered for biological relevance and compound availability, assessed for cytotoxicity, and validated by Hi-C. (**E**) Distribution of GCI gains and losses across drug-treated MCF7 samples. (**F)** Proportion of GCI containing different numbers of MCF7-specific enhancers at each anchor; other gene-related categories are shown in Fig. S11B. (**G**) Observed versus expected enrichment of cancer-associated SNPs overlapping both anchors in gene-gene (GG), gene-enhancer (GE), gene-other (GO), non-gene (NG), and intragenic (IG) interactions. (**H**) Reactome pathway enrichment of genes whose TSS or gene regions overlap anchors of GCIs colocalizing with enhancers at zero, one, or both anchors. Pathway names include their relative ranks by *p*-value. (**I-J**) Log2FC (library-normalized) changes in interaction strength for each loop category. (I) Loop types stratified by genomic annotation (e.g., gene-gene, intron-gene, exon-5′). Eper: eperezolid; Rade: radezolid. (J) Loop types stratified by enhancer colocalization in MCF7 cells treated with eperezolid or radezolid. (**K**) Reactome pathway enrichment of genes overlapping GCI anchors in DMSO-treated MCF7 cells, compared with those associated with GCI specifically lost (Log2FC ≤ -1) after eperezolid or radezolid treatment.

We next focused on MCF7 cells, which exhibited a balanced perturbational landscape and extensive compound coverage. Ranking compounds by predicted reductions in GCI identified eperezolid, radezolid, and JQ1 among the top candidates (**Fig. 6D-E, Fig. S11A**). To experimentally validate these predictions, we performed deep Hi-C profiling in MCF7 cells following short-term treatment. To ensure conceptual consistency with the GCI framework while leveraging the resolution of Hi-C data, we quantified treatment effects across anchor-defined chromatin interaction classes, including gene-associated interactions. Notably, eperezolid and radezolid induced pronounced reductions in gene-associated chromatin interactions, with the strongest effects observed in gene-gene interaction classes, whereas JQ1 showed a comparatively weaker and less specific impact under the same conditions (**Fig. 6I-J, Fig. S11D**). These results indicate that AI4Loop can prioritize compounds that selectively perturb specific structural components of the chromatin interaction landscape.

To further interpret these effects, we characterized the baseline chromatin interaction architecture in untreated MCF7 cells. Gene anchors showed strong enrichment for MCF7-specific enhancers and cancer-associated risk SNPs, and gene-associated interactions—particularly gene-gene loops—were preferentially linked to regulatory programs central to breast cancer biology, including chromatin remodeling, VEGF signaling, and estrogen signaling pathways (**Fig. 6F-H**, **Fig. S11B-C**). Consistently, gene-associated interactions preferentially disrupted by eperezolid and radezolid were enriched in estrogen- and cholesterol-related pathways (**Fig. 6K**), suggesting that these compounds target transcriptionally active regulatory hubs. Collectively, these results demonstrate that AI4Loop enables systematic identification of compounds that modulate oncogenic chromatin interaction programs and highlight gene-associated chromatin interactions as functionally relevant and drug-sensitive structural units in cancer.

## Discussion

The spatial organization of the genome plays a central role in regulating transcriptional programs by bringing regulatory elements and gene promoters into physical proximity within the nucleus (28,29). While technologies such as Hi-C and related chromosome conformation capture approaches have enabled direct mapping of chromatin architecture, their application across large biological cohorts remains limited by experimental cost and technical complexity (5,30). In parallel, DNA sequence-based models such as DeepC(4), Akita(31) and Orca(32) have shown that sequence features can robustly predict aspects of chromatin folding potential, highlighting the strong contribution of genome-encoded information to 3D genome organization. However, when applied to a common reference genome, DNA sequence-based models primarily capture static sequence-encoded folding potential; without sample-specific genomic, structural-variant, or epigenomic information, they are less suited to modeling dynamic chromatin interaction changes across tumors, tissues, or drug perturbations (33).

AI4Loop was therefore designed from a different perspective: to infer chromatin interactions directly from RNA-seq, an inexpensive, widely available and information-rich molecular readout. This design enables scalable prediction of sample-specific gene-centered chromatin interactions across large clinical and perturbational transcriptomic datasets, where matched Hi-C, HiChIP or whole-genome profiling is usually unavailable. In this study we demonstrate that transcriptomic profiles contain higher order genome organization information and can be used to infer chromatin interactions at scale. Using this principle, we developed AI4Loop, a deep learning framework that identifies GCI from RNA-seq data and enables systematic analysis of chromatin interaction dynamics across large transcriptomic datasets.

Our results suggested that GCI may contribute to malignant transcriptional regulation by coupling multiple genes within shared regulatory programs. Predicted interactions are enriched for active chromatin marks, transcriptional regulators, and RNA polymerase II occupancy, consistent with the known association between chromatin interactions and transcriptional activity (34). Moreover, the concordance of AI4Loop predictions with orthogonal chromatin conformation assays such as ChIA PET and Hi-TrAC, as well as its successful generalization to clinical samples, indicates that transcriptome-derived features capture aspects of regulatory chromatin architecture that extend beyond individual experimental platforms and controlled cell-line systems.

A central biological finding of this study is the pervasive increase in GCI across human cancers. Across more than 12,000 transcriptomes spanning 32 cancer types, most tumors exhibited elevated GCI relative to healthy tissues, particularly at oncogene-associated loci. Previous studies have shown that disruption of chromatin topology and enhancer rewiring can activate oncogenes during tumorigenesis (35,36). Our results suggest that, in addition to enhancer-promoter interactions, GCIs may also contribute to oncogenic transcriptional programs by forming multi gene regulatory hubs that reinforce coordinated gene expression. Such hubs may facilitate cooperative transcriptional activation among oncogenic pathways, thereby promoting tumor growth and survival.

Interestingly, GCI networks inferred by AI4Loop provided greater discriminatory power for cancer classification than gene expression alone. This observation suggests that the spatial coordination of transcriptional programs captured by chromatin interaction networks provides additional structure beyond individual gene expression levels. From a systems perspective, GCIs may therefore represent a higher order regulatory layer that integrates transcriptional activity across multiple genes. These findings are consistent with the idea that transcriptional programs emerge from coordinated regulatory activity within the 3D genome.

The study further demonstrates the value of transcriptome-inferred chromatin interactions for identifying therapeutic vulnerabilities. By integrating AI4Loop predictions with perturbational transcriptomic data from the LINCS Connectivity Map, we generated a large-scale atlas of chromatin interaction responses to drug treatments. This analysis identified candidate compounds predicted to reverse cancer associated GCI patterns. Experimental Hi-C profiling confirmed that the oxazolidinone antibiotics eperezolid and radezolid preferentially lost GCIs in breast cancer cells. Although these compounds are primarily known as antibacterial agents, our results suggest that they may also influence chromatin organization and transcriptional regulation in cancer cells. More broadly, these findings highlight the possibility of identifying chromatin modulating therapeutics through large scale transcriptomic perturbation datasets.

In the future, the simplicity of AI4Loop holds immense potential to expand its application by shifting its focus from GCI to different types of chromatin interactions. First, the current framework focuses on gene-centered chromatin interactions, but future work could expand it to additional classes of chromatin interactions, including inter-chromosomal and enhancer-associated interactions. Such extensions would enable broader investigation of long-range regulatory dynamics in cancer and during therapeutic perturbation. Second, AI4Loop can be readily applied to single-cell RNA-seq data through pseudo-bulk aggregation, providing an immediate route to study chromatin interaction programs in heterogeneous systems (**Fig. S12**). Longer term, dedicated model development may enable true single-cell-resolution inference, allowing direct analysis of cell-to-cell variability in 3D genome regulation. Third, integrating alternative RNA modalities, such as enhancer RNA or nascent RNA sequencing, may improve sensitivity to early regulatory events and extend inference toward enhancer-linked or co-transcriptional chromatin interaction programs (37).

In summary, AI4Loop provides a scalable framework for inferring chromatin interaction landscapes from widely available transcriptomic data. Our results reveal a pervasive gain of GCIs in cancer, show that these interaction profiles provide a structured representation of cancer state beyond expression alone, and demonstrate that transcriptome-inferred chromatin interaction programs can be exploited to prioritize compounds that perturb cancer-associated regulatory architecture. Together, these findings highlight the potential of integrating transcriptomic and structural genome information to uncover new principles of gene regulation and guide interaction-informed therapeutic strategies

## Materials and Methods

### AI4Loop benchmark dataset construction

To construct benchmark datasets for gene-centered chromatin interaction (GCI) prediction, we integrated Hi-C loop calls, RNA-seq profiles, CTCF ChIP-seq peaks, and gene annotations from four human cell lines, including K562, GM12878, HeLaS3, and IMR90 (**Fig. S13**). Hi-C loop calls were obtained from the high-resolution Hi-C datasets of Rao et al., in which loops were originally identified using the HiCCUPS pipeline implemented in Juicer. RNA-seq and CTCF ChIP-seq data were obtained from ENCODE, and gene annotations were downloaded from GENCODE Release 36 based on the GRCh37 genome assembly.

A GCI was defined as an intrachromosomal chromatin loop in which both loop anchors overlapped annotated gene regions. To reduce redundancy, anchors located within 500 bp were merged before interaction re-clustering. Loops with both anchors overlapping gene loci were retained as positive GCIs. Candidate interactions were restricted to gene-gene pairs within a linear genomic distance of 5 kb to 2 Mb, consistent with the distance range used for model training and downstream applications.

Negative samples were generated from four complementary sources to improve biological plausibility and reduce label noise. First, graph-based non-connected anchors were identified by constructing an interaction graph in which merged anchors were treated as nodes and Hi-C interactions as edges. Pairs of gene-overlapping anchors with no direct or indirect connection in the graph were defined as negative samples. Second, random CTCF-gene pairs were generated by sampling CTCF peaks from the same cell type, intersecting them with gene loci, resizing them according to the positive anchor-length distribution, and retaining pairs separated by 5 kb to 2 Mb that were not connected in the interaction graph. Third, random gene-gene pairs were sampled from the same cell line, required to satisfy the same distance range, resized to match the positive anchor-length distribution, and filtered to remove pairs connected by any Hi-C interaction or CTCF binding. Fourth, additional random gene pairs were sampled to match the genomic distance distribution of positive interactions, while excluding pairs overlapping Hi-C interaction anchors or CTCF peaks. The final negative set was assembled by aggregating non-overlapping samples from these four sources, with an approximate positive-to-negative ratio of 1:5.

### Multi-scale RNA-seq feature encoding

To represent transcriptional information around each candidate interaction anchor, we developed a multi-scale gene expression encoding strategy based on RNA-seq data. Each anchor was symmetrically extended by 30 kb upstream and downstream. The extended region was then partitioned into non-overlapping bins using seven spatial resolutions ranging from 1 kb to 7 kb. For each bin, the mean FPKM value of all genes overlapping that bin was calculated. Bins without overlapping genes were assigned a value of zero For each anchor, expression vectors generated at different binning resolutions were concatenated to form a unified multi-scale expression representation. For each candidate chromatin loop, the feature vectors from the left and right anchors were concatenated to represent the loop-level transcriptomic context. This design allowed AI4Loop to capture both local and broader transcriptional patterns surrounding candidate interacting gene loci.

### AI4Loop architecture and model training

AI4Loop uses a dual-branch bidirectional long short-term memory (Bi-LSTM) architecture to infer GCIs from RNA-seq-derived features. The multi-scale expression features from the two anchors of a candidate interaction were encoded separately by Bi-LSTM layers. The encoded representations from the two branches were then concatenated and passed through fully connected layers. A final sigmoid output layer generated the predicted interaction probability for each candidate gene pair.

Model performance was evaluated using two complementary splitting strategies. In the random-split validation, chromatin interaction samples were randomly divided into training and testing sets using an 80:20 ratio. This strategy assessed within-distribution model performance. In the chromosome-split validation, all samples involving chromosomes 4, 7, 8, and 11 were reserved as the test set, whereas samples from the remaining chromosomes were used for training. This strategy was used to evaluate model generalizability to unseen chromosomal contexts and to reduce potential inflation caused by local genomic dependencies.

Model performance was quantified using AUROC, AUPRC, accuracy, F1-score, and Matthews correlation coefficient. To benchmark AI4Loop against alternative architectures, we compared it with logistic regression, feedforward neural network, convolutional neural network, recurrent neural network, LSTM, and Transformer models. All models were trained and evaluated on the same datasets and train-test splits. Unless otherwise specified, neural network models were trained using the Adam optimizer, binary cross-entropy loss, a batch size of 30, and 30 training epochs.

### Pretrained model selection for downstream applications

AI4Loop models were trained separately using benchmark datasets from K562, GM12878, HeLaS3, and IMR90. For downstream applications, pretrained models were selected according to biological context rather than downstream outcome. The K562-pretrained model was used for leukemia-focused analyses because K562 is a hematopoietic cancer-derived cell line and is therefore most relevant to AML-related applications. The GM12878-pretrained model was used as the default reference model for pan-cancer TCGA analysis and drug perturbation screening because GM12878 provides a deeply characterized lymphoblastoid reference context and avoids anchoring pan-cancer inference to a single tumor lineage. For clinical CLL validation, both K562- and GM12878-pretrained models were evaluated because CLL is a hematopoietic malignancy and both models provide biologically relevant reference contexts. HeLaS3- and IMR90-trained models were primarily used for benchmarking, cross-cell generalization, and independent validation analyses.

### Cell-type-specific GCI identification and cross-cell evaluation

To assess whether AI4Loop could detect cell-type-specific interactions, chromatin interaction datasets from K562, GM12878, HeLaS3, and IMR90 were processed to identify GCIs unique to each cell type. Loop anchors were harmonized using a 10 kb positional tolerance to account for minor coordinate shifts between datasets. For each cell type, loops were defined as cell-type-specific if they were absent from the other three cell types. To control for genomic distance bias, distance-matched negative datasets were generated by pairing each cell-type-specific loop with a non-interacting gene pair of comparable genomic separation.

Pretrained AI4Loop models were then applied to predict interaction status in the cell-type-specific datasets. Prediction performance was evaluated using precision-recall curves, AUROC, and AUPRC. This analysis was designed to test whether AI4Loop could generalize beyond shared loops and detect interactions specific to individual cellular contexts.

### Evaluation using clinical and orthogonal chromatin interaction datasets

To evaluate generalizability beyond training cell lines and Hi-C-derived loop calls, AI4Loop was tested on clinical samples and independent chromatin interaction datasets generated by orthogonal technologies. For clinical validation, AI4Loop was applied to six chronic lymphocytic leukemia (CLL) samples with matched RNA-seq and Hi-C data. Given the hematopoietic origin of CLL, both GM12878- and K562-pretrained models were used to predict chromatin interactions in these patient samples.

For platform-independent validation, AI4Loop was evaluated on Hi-TrAC and ChIA-PET datasets. Raw Hi-TrAC data were obtained from GEO under accession number GSE1801755(20), and ChIA-PET data were retrieved from GSE728166(38). Because AI4Loop was designed to predict gene-centered chromatin interactions, analyses of these datasets were restricted to TSS-associated loops, which represent gene-centric 3D interactions. For each candidate loop, AI4Loop generated a prediction probability between 0 and 1. Candidate loops were ranked by predicted probability, and the top and bottom ranked loops were used as high-confidence and low-confidence groups for downstream comparison of experimental chromatin interaction and epigenomic signals.

### AI4Loop performance validation in Acute Myeloid Leukemia samples

To identify AML-associated GCIs, we used the K562-pretrained AI4Loop model, selected a priori because K562 is a hematopoietic cancer-derived cell line. All intrachromosomal gene pairs within 5 kb to 2 Mb were generated, yielding 3,590,596 candidate pairs from 61,860 annotated genes. Gene pairs were ranked according to their predicted interaction probabilities, and the top and bottom 1,000 pairs were defined as high-confidence and low-confidence predicted GCIs, respectively.

RNA-seq data from 40 AML and 38 healthy samples in the TARGET-AML cohort (39) were then encoded using the top 1,000 high-confidence K562-derived GCI features, generating a 1,000 × 78 probability matrix. AML-associated GCIs were identified by comparing the average predicted probabilities between AML and healthy samples using a probability threshold of 0.5. Transcription factor binding and histone-mark annotations used for functional interpretation were obtained from ENCODE (**Table S1**) (40).

### AI4Loop application to TCGA pan-cancer transcriptomes

For pan-cancer analysis, we used the GM12878-pretrained AI4Loop model as the default reference model to predict GCIs across TCGA samples. A total of 12,347 TCGA RNA-seq samples spanning 32 cancer types and matched normal tissues were collected. Gene expression profiles included 55,420 annotated genes. Genes with zero expression across all samples were first removed. We then calculated the average FPKM value of each gene across cancer types and retained genes with an average FPKM greater than 5, resulting in 5,338 commonly expressed genes with robust transcriptional activity.

Candidate gene pairs were restricted to intrachromosomal pairs located 5 kb to 2 Mb apart, yielding 49,255 potential gene pairs. For each TCGA sample, AI4Loop predicted the interaction probability of each candidate gene pair. Gene pairs with prediction probability greater than 0.5 were classified as predicted GCIs. Based on their prevalence across cancer and normal samples, predicted GCIs were further grouped into shared GCIs (GCI-all), cancer-enriched GCIs (GCI-cancer), healthy-enriched GCIs (GCI-healthy), cancer-type-specific GCIs (GCI-cancer-one), and heterogeneous GCIs (GCI-several) detected across multiple but not all sample groups. For each sample, the total number of predicted GCIs was calculated and compared with the corresponding average gene expression level to assess the relationship between inferred chromatin interaction burden and transcriptional activity.

### Comparison of GCI, expression, and co-expression representations

To evaluate whether AI4Loop-predicted GCIs provide information beyond gene expression, we compared GCI-based sample representations with matched gene expression and co-expression representations. These comparisons were performed using cancer-type-specific GCIs, cancer-enriched and healthy-enriched GCIs, and landmark gene-derived GCIs. UMAP was used for visualization, and clustering performance was quantified using adjusted Rand index, normalized mutual information, silhouette coefficient, Davies-Bouldin index, and Calinski-Harabasz score.

### Drug perturbation analysis using CMap L1000 profiles

To identify compounds predicted to modulate GCIs, AI4Loop was applied to drug-induced transcriptomic profiles from the LINCS Connectivity Map L1000 platform, including Phase I and Phase II datasets(10,41). Because the L1000 platform directly measures 978 landmark genes, drug perturbation analysis was restricted to landmark gene pairs located on the same chromosome and separated by 5 kb to 2 Mb. This filtering yielded 1,290 candidate landmark gene pairs.

For drug perturbation screening, the GM12878-pretrained AI4Loop model was used as the default reference model to provide a common model across diverse cell lines and perturbation conditions. Because L1000 profiles are represented by transcriptional activity scores rather than RNA-seq FPKM values, transcriptional activity scores were normalized to the GM12878 FPKM distribution using a cumulative distribution matching strategy. The normalized values were then used as AI4Loop input.

For each perturbation profile, AI4Loop generated predicted GCI probabilities for all landmark gene pairs. Interactions with prediction probability greater than 0.5 were defined as confident GCIs. Compounds were ranked according to their predicted ability to induce GCI gain or loss relative to the corresponding control condition. The resulting perturbation-GCI matrix was used to identify candidate compounds predicted to modulate cancer-associated chromatin interaction programs.

### Hi-C validation of drug-induced chromatin interaction changes

MCF7 cells were treated with selected compounds and subjected to cell viability assessment and Hi-C profiling. Cell viability was assessed after 72 h treatment using crystal violet staining. Cells were fixed with 4% glutaraldehyde, stained with 0.1% crystal violet in 10% ethanol, washed, air-dried, and lysed in 10% acetic acid. Absorbance was measured at 592 nm to estimate relative cell density.

Hi-C libraries were generated using the Dovetail TopoLink V2 kit according to the manufacturer’s protocol and sequenced on the Illumina NovaSeq 6000 platform with 2 × 150 bp paired-end reads. Raw sequencing reads were trimmed using Trimmomatic and aligned to the human reference genome hg19 using BWA (42,43). Pairtools was used to parse and deduplicate aligned read pairs, followed by sorting and indexing using samtools and pairix. Contact matrices were generated using cooler and balanced using the built-in balancing option.

Hi-C cool files were converted to sparse interaction matrices at 4 kb resolution. Only intrachromosomal contacts were retained. Interactions with fewer than two counts and interactions shorter than 4 kb were excluded. Sparse matrices from all samples were merged into a unified matrix. Individual interaction sites were extracted as anchors and annotated using ChIPSeeker. Cancer-associated SNPs were obtained from the GWAS Catalog by filtering traits containing the term “cancer.” MCF7-specific enhancer annotations were obtained from the UCSC EPInteract database.

For each treatment, library-normalized interaction counts were calculated and stratified by interaction type. Treatment-control ratios were computed relative to DMSO after adding a pseudocount of 0.1 to all values, followed by log2 transformation. Interaction classes with sufficient numbers of interactions were retained for downstream comparison. Reactome pathway enrichment analysis was performed using ReactomePA based on genes associated with promoter- or TSS-containing interaction classes.

### Application of AI4Loop to pseudo-bulk single-cell RNA-seq perturbation profiles

To explore whether AI4Loop could be extended to single-cell perturbation data, publicly available single-cell RNA-seq drug perturbation datasets were analyzed. Cells were grouped by cell line and treatment condition, and expression profiles were aggregated into pseudo-bulk profiles. Pseudo-bulk gene counts were converted to FPKM values using gene length annotations from the hg19 reference genome. AI4Loop was then applied directly to the pseudo-bulk FPKM profiles without model retraining. Predicted GCIs were defined using a probability threshold of 0.5, consistent with the TCGA and drug perturbation analyses.

## Data and code availability

All datasets used in this study are publicly available or generated as part of this work. The pan-cancer chromatin interaction atlas generated by AI4Loop has been deposited in the Nanyang Technological University data repository (https://doi.org/10.21979/N9/ORBU74). Drug-perturbed Hi-C datasets generated in this study are available through GEO under accession number GSE287383. The AI4Loop code, trained models, and analysis scripts are available at https://github.com/DaoFuying/AI4Loop.

## Acknowledgments

We thank members of the Fullwood laboratory for discussion and advice. We acknowledge the Eric and Wendy Schmidt AI in Science Postdoctoral Fellowship (F.D and B.L) and NTU AI4X Postdoctoral Fellowship (F.D). This research is supported by the National Research Foundation, Singapore under its AI Singapore Programme (AISG Award No: AISG3-GV-2023-014) and by the Ministry of Education, Singapore, under its Academic Research Fund Tier 1 (RG38/23), both awarded to M.J.F. (PI) and K.C.K. (Co-I). This research is also supported by the Singapore Ministry of Health’s National Medical Research Council under its Singapore Translational Research Investigator Award STaR (MOH-000709) awarded to G.B.C (PI) and M.J.F(Co-I), and the National Research Foundation, Singapore (NRF-PA2025-NTU; Award No. 026374-00004) awarded to F.D (PI).

## Supporting Information

Supplementary Methods and Figs. S1-S13 are provided in the Supporting Information. Tables S1-S3 are available at the AI4Loop GitHub repository (https://github.com/DaoFuying/AI4Loop).

## Author Contributions

F.D. performed formal analysis, software development, visualization, conceptualization, investigation, methodology, data curation, validation, and writing. B.L. contributed conceptualization, methodology, validation, formal analysis, investigation, visualization, supervision, and writing. P.A. performed formal analysis and investigation. W.K.T. and X.L. contributed to methodology, investigation and software. B.C.G. and W.J.C. contributed to writing review and editing, and B.C.G. also acquired funding. C.K.K., H.Lin, and H.Lyu contributed to methodology and writing; C.K.K. and H.Lyu also provided supervision. M.J.F. contributed conceptualization, investigation, writing, funding acquisition, supervision, and project administration.

## Competing Interest Statement

The authors declare no competing interests.

## Notes

https://doi.org/10.21979/N9/ORBU74

https://github.com/DaoFuying/AI4Loop

https://www.ncbi.nlm.nih.gov/geo/query/acc.cgi?acc=GSE287383

### Competing Interest Statement

The authors have declared no competing interest.

