## Supplementary Information for "AI4Loop: an Artificial Intelligence Framework Reveals Increased 3D Chromatin Interactions and Therapeutic Vulnerabilities across 12,000 Cancer Samples"

**The PDF file includes:**

Supplementary Methods
Figs. S1 to S13
References

### Supplementary Methods

#### Public datasets and genome annotations

To construct AI4Loop and evaluate its generalizability, we integrated public Hi-C, RNA-seq, CTCF ChIP-seq, ChIA-PET, Hi-TrAC, cancer transcriptome, drug perturbation, and single-cell RNA-seq datasets. Hi-C loop calls for K562, GM12878, HeLaS3, and IMR90 were obtained from the high-resolution Hi-C datasets reported by Rao et al[1]. These loops were originally identified using HiCCUPS implemented in the Juicer suite. HiCCUPS detects statistically significant chromatin loops by identifying local enrichment of Hi-C contact frequencies relative to local background neighborhoods, followed by false discovery rate control. These loop calls were used as the reference positive chromatin interaction set for AI4Loop benchmark construction.

Matched RNA-seq and CTCF ChIP-seq datasets for K562, GM12878, HeLaS3, and IMR90 were obtained from ENCODE[2]. Human gene annotations were downloaded from GENCODE Release 36 based on the GRCh37/hg19 genome assembly for model construction and benchmark generation. For newly generated Hi-C data in MCF7 cells, sequencing reads were aligned to hg19, as described below. All genomic analyses were performed using consistent genome assemblies within each analysis module, and genome versions are specified for each dataset where applicable.

#### Definition of gene-centered chromatin interactions

AI4Loop was designed to infer gene-centered chromatin interactions (GCIs) from RNA-seq data. In this study, a GCI was defined as an intrachromosomal chromatin interaction in which both anchors overlapped annotated gene regions. We focused on gene-gene interactions because AI4Loop uses gene expression signals as input and therefore requires candidate loci with measurable transcriptomic features. Interchromosomal pairs were excluded from model training and downstream applications. Candidate gene pairs were restricted to a linear genomic distance of 5 kb to 2 Mb, consistent with the range in which high-confidence focal chromatin interactions are commonly evaluated from available loop calls and to reduce the inclusion of very long-distance contacts with limited loop-calling reliability.

#### Positive sample construction

To construct positive samples, Hi-C loop anchors were first intersected with annotated gene regions. Anchors overlapping gene loci were retained (**Fig. S13**). To reduce redundancy caused by nearby loop anchors, anchors located within 500 bp were merged. Chromatin interactions were then re-clustered using these merged anchors. A loop was retained as a positive GCI if both anchors overlapped annotated gene regions after anchor merging. When multiple genes overlapped the same anchor, all possible gene-pair instances were generated and subsequently collapsed to unique gene pairs where necessary to avoid redundant entries. All positive samples were restricted to intrachromosomal gene-gene pairs separated by 5 kb to 2 Mb.

#### Negative sample construction

Negative samples were generated from four complementary sources to improve biological plausibility, reduce label noise, and control for potential biases associated with genomic distance and CTCF occupancy (**Fig. S13A**):

First, graph-based non-connected anchors were generated from the Hi-C interaction network. In this network, each merged anchor was treated as a node and each Hi-C interaction was treated as an edge. Pairs of gene-overlapping anchors that were neither directly nor indirectly connected in the graph, meaning that no path existed between them, were defined as negative samples.

Second, random CTCF-gene pairs were generated. CTCF peaks from the same cell type were intersected with gene loci. For each such region, an anchor length was sampled from a normal distribution fitted to the anchor-size distribution of positive samples. If required, the region was symmetrically extended to match the sampled anchor size. All pairs of such regions separated by 5 kb to 2 Mb and not connected in the interaction graph were retained as negative samples.

Third, random gene-gene pairs were sampled from the same cell line. These pairs were required to satisfy three criteria: both genes resided within a genomic distance of 5 kb to 2 Mb; gene regions were resized to match the anchor-length distribution of positive samples; and the gene pair was not connected through any known Hi-C interaction or CTCF binding. To reduce redundancy, gene-gene random pairs overlapping with pairs already included in the CTCF-gene random set were removed.

Fourth, distance-matched random gene pairs were generated to control for genomic distance effects[3]. Random gene pairs were sampled to match the genomic distance distribution of positive interactions. Pairs overlapping Hi-C interaction anchors or CTCF peaks were excluded.

The final negative dataset was assembled by aggregating all non-overlapping samples from these four sources. A target positive-to-negative ratio of approximately 1:5 was used for model training and evaluation. All positive and negative samples were restricted to gene-gene pairs to ensure consistency with the prediction task of AI4Loop.

**Distance matching and benchmark dataset statistics**

Because genomic distance is a major determinant of chromatin contact frequency, distance matching was performed to reduce distance-driven performance inflation. For each cell line, the genomic distance distribution of negative samples was matched to that of positive samples within the 5 kb to 2 Mb range. The distance distributions of positive and negative samples were compared using the Kolmogorov-Smirnov test. The numbers of positive and negative samples, chromosome-wise distributions, and distance distributions for K562, GM12878, HeLaS3, and IMR90 are shown in **Fig. S13**.

For performance evaluation, AUROC and AUPRC were calculated using held-out positive and negative samples. AUPRC values were interpreted relative to the positive-sample prevalence in the corresponding test set, which represents the random baseline under class imbalance. Unless otherwise specified, predicted probabilities were used directly for AUROC and AUPRC calculation without applying a fixed classification threshold.

**Multi-scale gene expression features extraction from RNA-Seq data**

To extract informative transcriptional features surrounding chromatin anchors, we implemented a multi-scale gene expression (GE) encoding scheme based on RNA-Seq data. Each anchor was symmetrically extended by 30 kb upstream and downstream, forming a region of total length $L$.

To capture expression variation across different spatial resolutions, we applied seven binning strategies with bin sizes $bin_b \in \{1kb,\ 2kb, \cdots, 7kb\}$. As shown in Eq. (1), each extended anchor region was divided into $n_b = L/bin_b$ non-overlapping bins for binning scale *b*. For each bin *i*, the expression value $G_{bi}$ was calculated as the mean FPKM of all genes overlapping that bin; if no gene overlapped, the value was set to zero as defined in Eq. (3). The resulting expression profile at scale *b* was represented as a feature vector $G_b$, as shown in Eq. (2).

Finally, all scale-specific vectors were concatenated to form a unified multi-scale expression feature vector *GE* (see Eq. (1)). For each chromatin loop, feature vectors from the left and right anchors were concatenated to represent the full loop-level expression context. This design captures both local fine-grained and broad-scale transcriptional signals, enabling robust inference of spatial gene-gene regulatory relationships directly from RNA-Seq data.

For chromatin loops, features from the left and right anchors were concatenated to yield the final loop-level expression feature. This strategy captures both fine-resolution and broader transcriptional patterns, facilitating robust learning of spatial gene-gene regulatory relationships from RNA-Seq alone.

$$GE = \bigcup_{b=1}^{7} G_b \tag{1}$$

$$G_b = \left[G_{b1}, G_{b2}, \cdots, G_{bn_b}\right], n_b = \frac{L}{bin_b} \tag{2}$$

$$G_{bi} = \begin{cases} \frac{1}{k}\sum_{j=1}^{k} FPKM_j\ , if\ i\ overlaps\ with\ k\ genes \\ 0, if\ no\ genes\ overlaps\ with\ bin\ i \end{cases} \tag{3}$$

**AI4Loop framework**

Bi-directional Long Short-Term Memory (Bi-LSTM) networks, a type of Recurrent Neural Network (RNN), have shown superior performance in various tasks over traditional unidirectional LSTMs and RNNs due to their ability to process data from both past and future states simultaneously [4]. As shown in **Fig. 1B**, Bi-LSTM layer with 16 units is employed to encode the gene expression feature set from both anchors separately. This approach allows the model to consider the expression patterns from both anchors, ensuring a comprehensive understanding of the gene interactions within a chromatin loop. After encoding, the output vectors from the Bi-LSTM layers are concatenated and fed into fully connected layers. This sequential processing facilitates the capture of complex patterns in gene expression data, leveraging the strengths of deep learning to analyze chromatin structures. In last dense layer in the model uses the sigmoid activation function to generate classification probability for a query chromatin loop.

**Model training and evaluation strategies**

To ensure a comprehensive and robust assessment of model performance, we employed two complementary data splitting strategies for training and evaluation: random-split validation and chromosome-split validation. These two approaches differ in how training and testing data are partitioned, offering insights into model generalization under different biological assumptions.

Random-split validation. In the random-split strategy, chromatin interaction samples (i.e., gene-gene pairs with associated features and labels) were randomly partitioned into training and testing subsets. Specifically, 80% of the samples were randomly selected to form the training set, while the remaining 20% were used for testing. This approach ensures that training and test samples are drawn from the same global distribution and may include anchors located on the same chromosomes. As such, this strategy primarily evaluates the model's ability to fit and generalize within a similar sample space, but does not assess cross-chromosomal or spatial generalizability.

Chromosome-split validation. To test the model's capacity for generalizing to unseen chromosomal contexts, we adopted a chromosome-based splitting strategy. In this setting, all GCI samples involving chromosomes 4, 7, 8, and 11 were reserved exclusively as the test set. The remaining chromosomes were used to generate the training set. This design ensures that none of the test anchors appear in the training phase, providing a stringent test of the model's ability to generalize to entirely new chromosomal regions. This approach more closely mimics real-world applications where the model may be deployed on new chromosomes, cell types, or patient samples.

By comparing model performance across these two strategies, we evaluate not only within-distribution accuracy but also the biological transferability of the model to unseen genomic contexts.

**Benchmarking against alternative architectures**

To systematically benchmark our Bi-LSTM–based AI4Loop framework, we compared it with six representative baseline architectures, including both classical machine learning and deep learning models (**Fig. 2I-J**, **Fig. S5**). All models were trained and evaluated on the same datasets, with an 80/20 train–test split applied consistently across all methods to ensure fairness. Unless otherwise specified, training used the Adam optimizer (learning rate = 0.001), batch size = 30, 30 epochs, and binary cross-entropy loss. Model performance was evaluated using AUROC, AUPRC, ACC, MCC, and F1.

Logistic Regression (LR). Logistic regression was implemented in scikit-learn. Input features were standardized with StandardScaler. The model employed L2 regularization with the liblinear solver, class_weight="balanced" to address class imbalance, a maximum of 5000 iterations, and random_state=42 for reproducibility.

Feedforward Neural Network (FFNN). In the FFNN baseline, gene expression features from the left and right anchors were flattened and concatenated. The merged feature vector was passed through fully connected layers with ReLU activation and optional dropout regularization (dropout rate = 0.3). The final output layer consisted of a single sigmoid unit predicting loop probability. A typical architecture included two hidden layers with 64 and 32 units.

Convolutional Neural Network (CNN). The CNN baseline encoded left and right anchor features separately using consecutive 1D convolutional layers (kernel size = 4, 64 filters), each followed by LeakyReLU activation ($\alpha = 0.128$) and MaxPooling1D (pool size = 2). The resulting feature maps were flattened, concatenated, and passed through fully connected layers with ReLU activation and dropout. A final sigmoid layer produced the interaction probability.

Recurrent Neural Network (RNN). The RNN baseline was implemented using gated recurrent units (GRU). Left and right anchor features were independently processed by a GRU layer (64 units, return sequences enabled). The outputs were flattened and concatenated, followed by one or more fully connected layers with ReLU activation and optional dropout. The final output layer was a sigmoid unit.

Long Short-Term Memory (LSTM). The LSTM baseline used the same structure as the GRU-based RNN, but with LSTM units (64 hidden units, return sequences enabled). Flattened outputs from left and right anchors were concatenated and passed through dense layers with ReLU activation and dropout, followed by a sigmoid output layer.

Transformer. The Transformer baseline concatenated left and right anchor features along the sequence axis and processed them with two stacked Transformer encoder blocks. Each block consisted of multi-head self-attention (key dimension = 64, 4 heads, dropout = 0.1), residual connections, LayerNormalization, and a position-wise feed-forward network (Dense 128 → Dense projection). The final sequence representation was flattened and passed through a multilayer perceptron head (Dense 128 → Dense 64, ReLU, dropout = 0.1). A single sigmoid unit produced the loop probability.

**Ablation and feature-design analyses**

To evaluate the contribution of different feature designs, we performed a series of ablation analyses. First, we compared models trained with gene expression features derived from short genes, long genes, cancer-associated genes, randomly selected genes, highly expressed genes, enhancer-associated genes, and silencer-associated genes. Enhancer-associated genes were defined as genes colocalized with H3K27ac, whereas silencer-associated genes were defined as genes colocalized with H3K9me3.

Second, we compared window-based and non-window-based feature encoding strategies to evaluate the contribution of local transcriptional context around anchors. Third, we compared features extracted from anchors, the window region between anchors, and the full loop region. Fourth, we evaluated different window-based aggregation strategies, including mean, maximum, and variance. Fifth, we compared models trained using genomic distance alone, RNA-seq features alone, and the combination of genomic distance and RNA-seq features. These analyses were used to determine whether AI4Loop captured information beyond simple genomic distance and expression intensity.

**Feature attribution analysis**

To interpret AI4Loop predictions, feature attribution analyses were performed using SHAP. SHAP values were calculated for representative trained models to identify input features that contributed most strongly to interaction prediction. The top-ranked features were visualized using SHAP summary plots. For selected cell lines, volcano plots were generated to compare feature differences between positive and negative samples, with features ranked by mean absolute SHAP values and log2 fold changes.

To evaluate the effect of anchor window size, five-fold cross-validation was performed across different window sizes ranging from 1 kb to 7 kb. Precision-recall curves and AUPRC values were compared across window sizes to assess the stability of prediction performance.

**Pretrained model selection and downstream use**

AI4Loop models were trained separately using benchmark datasets from K562, GM12878, HeLaS3, and IMR90. For downstream applications, pretrained models were selected according to biological context rather than downstream outcome.

The K562-pretrained model was used for leukemia-focused analyses because K562 is a hematopoietic cancer-derived cell line and is therefore most relevant to AML-related applications. The GM12878-pretrained model was used as the default reference model for pan-cancer TCGA analysis and drug perturbation screening because GM12878 provides a deeply characterized lymphoblastoid reference context and avoids anchoring pan-cancer inference to a single tumor lineage. For clinical CLL validation, both K562- and GM12878-pretrained models were evaluated because CLL is a hematopoietic malignancy and both models provide biologically relevant reference contexts. HeLaS3- and IMR90-trained models were primarily used for benchmarking, cross-cell generalization, and independent validation analyses.

**Cell-type-specific GCI identification and evaluation**

Chromatin interaction datasets from K562, GM12878, HeLaS3, and IMR90 were processed to identify cell-type-specific GCIs. Loop anchors were harmonized using a 10 kb positional tolerance to ensure that minor coordinate shifts did not affect overlap detection. For each cell type, loops were defined as specific if they were absent from the other three cell types, and shared and unique interaction sets were quantified and visualized using a four-way Venn diagram (**Fig. S3A**). To control for genomic distance biases, distance-matched negative datasets were generated by pairing each specific loop with a non-interacting locus pair of comparable genomic separation, resulting in balanced positive and negative datasets for each cell type (**Fig. S3B-D**). Pre-trained AI4Loop models (independently trained on K562 and GM12878 data) were then applied to predict interaction status in the cell-type-specific datasets. Model performance was assessed using precision-recall curves, AUROC, and AUPRC metrics across all test cell types (**Fig. S3E-I**). This evaluation framework directly measures the ability of AI4Loop to detect chromatin interactions that are unique to individual cell types, thereby testing model generalizability beyond shared gene expression patterns.

**Evaluation on clinical CLL samples**

To assess model performance in clinical samples, AI4Loop was applied to six chronic lymphocytic leukemia samples with matched RNA-seq and Hi-C data[3]. Two samples, CLL-102 and CLL-344, belonged to the IGHV-unmutated subtype, whereas CLL-312, CLL-324, CLL-401, and CLL-484 belonged to the IGHV-mutated subtype. Because CLL is a hematopoietic malignancy, AI4Loop models trained on GM12878 and K562 were used for prediction.

For each CLL sample, candidate gene-centered interactions were generated and scored using AI4Loop. Predicted probabilities were compared with matched Hi-C interaction evidence. To further assess confidence-dependent concordance, candidate loops were ranked by predicted probability. High-confidence and low-confidence groups were defined from the top-ranked and bottom-ranked predictions, respectively, and compared with Hi-C contact intensity and aggregate peak analysis.

Distance-stratified prediction performance was further evaluated by grouping candidate interactions according to genomic distance. Correct prediction ratios were calculated within each distance bin to assess whether AI4Loop retained predictive power across different genomic distance ranges.

**Evaluation on Hi-TrAC and ChIA-PET datasets**

To evaluate platform independence, AI4Loop was tested on independent chromatin interaction datasets generated using Hi-TrAC and ChIA-PET. Raw Hi-TrAC data were obtained from GEO accession GSE1801755[5],, and ChIA-PET data were obtained from GEO accession GSE728166[6]. Because AI4Loop was designed to infer gene-centered interactions, we restricted the evaluation to TSS-associated loops that represent gene-centric 3D interactions.

For each candidate loop, AI4Loop generated a prediction probability between 0 and 1. Candidate loops were ranked by prediction probability. The top 1,000 predictions were defined as the high-confidence group, and the bottom 1,000 predictions were defined as the low-confidence group. These groups were compared with experimental interaction evidence and epigenomic annotations, including active histone marks, enhancer/promoter annotations, and transcription factor or RNA polymerase II occupancy.

**TCGA pan-cancer RNA-seq sample collection**

To investigate GCI landscapes across cancer types, AI4Loop was applied to RNA-seq profiles from The Cancer Genome Atlas (TCGA)[7]. A total of 12,347 RNA-seq samples spanning 32 cancer types and matched normal tissues were collected (Fig. 4A-C). The expression matrix contained 55,420 annotated genes.

To define a high-confidence set of candidate genes and reduce biologically uninformative combinations, genes with zero expression across all samples were first excluded. The average FPKM value of each gene was then calculated across cancer types. Genes with an average FPKM greater than 5 were retained, resulting in 5,338 commonly expressed genes with robust transcriptional activity (Fig. S9A). Candidate gene pairs were restricted to intrachromosomal pairs separated by 5 kb to 2 Mb, yielding 49,255 potential gene pairs for downstream prediction.

**TCGA GCI prediction and classification**

For each TCGA sample, AI4Loop was used to compute prediction probabilities for all 49,255 candidate gene pairs. The GM12878-pretrained model was used as the default reference model for this pan-cancer analysis. A gene pair was considered a predicted GCI in a given sample if its prediction probability exceeded 0.5.

Gene pairs with no predicted interaction signal across all samples were defined as non-GCIs. Gene pairs with predicted interaction signals in at least one sample were defined as candidate GCIs. Based on prediction patterns across cancer and normal samples, GCIs were grouped into five categories: GCI-all, representing gene pairs consistently predicted as interacting across cancer and normal samples; GCI-cancer, representing gene pairs predicted as interacting in cancer samples but not normal samples; GCI-healthy, representing gene pairs predicted as interacting in normal samples but not cancer samples; GCI-cancer-one, representing interactions detected specifically in one cancer type; and GCI-several, representing interactions detected across several but not all cancer types.

**Quantification of GCI counts and global expression level for TCGA samples**

To quantify the abundance of GCI across different cancer types, we first computed the number of predicted GCIs for each individual sample. Specifically, for each of the 49,255 candidate gene pairs, if the prediction probability generated by AI4Loop exceeded 0.5 in a given sample, the pair was considered an active interaction in that sample. The total number of such gene pairs was recorded as the GCI count for that sample. Subsequently, we averaged the GCI counts across all samples within the same cancer type to obtain the mean GCI burden for that cancer type.

To evaluate the global transcriptional activity of each sample type, we calculated the mean FPKM value of all retained genes (i.e., the 5,338 commonly expressed genes) within each individual sample. These per-sample average expression values were then averaged across all samples of a given cancer type, resulting in the mean gene expression level for that group. This metric serves as a representative measure of overall gene expression activity.

Together, these two metrics, GCI count and mean gene expression, provide complementary views of chromatin interaction dynamics and transcriptional output across tumor types, enabling comparative analyses of 3D genome regulatory complexity in cancer (**Fig. S12F-H**).

**Comparison of GCI, gene expression, and co-expression representations**

To determine whether AI4Loop-predicted GCI profiles provide information beyond gene expression, we compared three sample representations: AI4Loop-predicted GCI probabilities, expression levels of genes involved in the corresponding GCI sets, and co-expression features derived from the same gene pairs.

Comparisons were performed using three feature sets: cancer-type-specific GCIs, cancer-enriched and healthy-enriched GCIs, and landmark gene-derived GCIs. UMAP was used to visualize sample structure. Quantitative clustering performance was evaluated using adjusted Rand index, normalized mutual information, silhouette coefficient, Davies-Bouldin index, and Calinski-Harabasz score. Higher adjusted Rand index, normalized mutual information, silhouette coefficient, and Calinski-Harabasz score indicate better clustering performance, whereas lower Davies-Bouldin index indicates better compactness and separation.

**Motif enrichment analysis**

Motif enrichment analysis was performed for selected cancer-associated GCI sets. Genomic sequences corresponding to GCI anchors were extracted, and enriched transcription factor motifs were identified using standard motif enrichment procedures. Enriched motifs were ranked by statistical significance and visualized for pan-cancer and cancer-type-specific GCI sets.

**CMap L1000 perturbation data collection**

To identify compounds capable of modulating GCIs, AI4Loop was applied to large-scale drug-induced transcriptomic profiles from the Connectivity Map L1000 platform generated by the

LINCS consortium. L1000 data were collected from LINCS Phase I and Phase II datasets, corresponding to GSE92742 and GSE70138[8, 9]. These datasets contain drug-induced transcriptional responses across 42 human cell lines under different compound, concentration, and time-point conditions.

**Transcriptional Activity Score (TAS) normalization for AI4Loop input**

The L1000 platform directly measures the expression levels of 978 landmark genes, which are used to infer the transcriptome-wide response to perturbagens. Instead of raw expression values, L1000 profiles are typically represented by a summary metric known as the Transcriptional Activity Score (TAS). TAS is a composite measure that integrates signature strength (SS) and replicate correlation (CC) to quantify the overall transcriptional impact of a compound. A higher TAS indicates greater transcriptional perturbation of landmark genes in response to treatment.

To adapt L1000 data for input to AI4Loop, we developed a normalization strategy to align TAS values with the RNA-Seq-based expression profiles used in model training. Specifically, we employed the FPKM distribution from the GM12878 cell line as the reference baseline.

For a given set of TAS values $b = \{b_1, b_2, \cdots, b_n\}$ and a set of FPKM values $a = \{a_1, a_2, \cdots, a_m\}$, we can calculate the cumulative distribution function (CDF) of the reference and target distributions by Eq. (4), in which, the $I$ is the indicator function. Then mapping the target distribution $b$ to the range of the reference distribution $b$ in Eq. (5), in which $CDF_a^{-1}$ represents the inverse function of $CDF_a$, and used to align points in the target distribution $b$ with values in the reference distribution $b$. Finally, we can have the final normalized TAS values $b' = \{b'_1, b'_2, \cdots b'_n\}$.

$$CDF_a(x) = \frac{1}{m}\sum_{i=1}^{m} I(a_i \leq x),\ CDF_b(x) = \frac{1}{n}\sum_{j=1}^{n} I\left(b_j \leq x\right) \quad (4)$$

$$b'_j = CDF_a^{-1}\left( CDF_b\left(b_j\right)\right), j = 1, 2, \cdots, n \quad (5)$$

This normalization preserves the rank information encoded in the TAS vector while projecting it into the range of biologically observed expression values. The resulting $b'$ values are used as input to AI4Loop for chromatin interaction prediction.

**Landmark GCI predictions for screening drugs**

Given that the L1000 assay directly quantifies the expression of only 978 landmark genes, our chromatin interaction analysis was restricted to gene pairs within this predefined set. To ensure biological plausibility and maintain consistency with the genomic distance constraints used in model training, we considered all landmark gene pairs located on the same chromosome and separated by a linear genomic distance ranging from 5 kb to 2 Mb. This filtering yielded a total of 1,290 candidate gene-gene pairs for downstream evaluation.

Each of these gene pairs was then assessed using AI4Loop across all available perturbation profiles, enabling the systematic prediction of GCIs under diverse compound treatment conditions. To identify high-confidence interaction events, a prediction probability threshold of 0.5 was applied to binarize predicted interactions, providing a stringent criterion for distinguishing interaction gains or losses induced by drug perturbations. The resulting compound-interaction probability matrix captures predicted GCI modulation across thousands of small-molecule exposures and cell types. This matrix constitutes a high-resolution resource for identifying pharmacological agents that are predicted to induce or disrupt GCI contacts, offering mechanistic insights into compound action and informing opportunities for drug repurposing based on 3D genome architecture.

**Perform Hi-C experiments to verify drug function and analysis**

Cell Viability Assay for Hi-C experiments. To assess drug cytotoxicity and determine appropriate treatment doses, MCF7 cells were cultured in DMEM supplemented with 10% fetal bovine serum (FBS) and seeded into 24-well plates. Cells were treated with selected compounds for 72 hours at concentrations specified in the corresponding results section. After incubation, cell viability was assessed via crystal violet staining: cells were fixed with 4% glutaraldehyde, stained with 0.1% crystal violet in 10% ethanol, washed, air-dried, and lysed in 10% acetic acid. Absorbance was measured at 592 nm using a plate reader to quantify relative cell density.

Hi-C library preparation and sequencing. Hi-C experiments were performed using the Dovetail™ TopoLink™ V2 kit according to the manufacturer's standard protocol. Library construction was carried out by NovogeneAIT, and sequencing was performed on the Illumina NovaSeq 6000 platform with a read length of 2 × 150 bp, achieving a sequencing depth of 800 million paired-end reads per sample.

Hi-C data processing and contact map generation. Raw sequencing reads were preprocessed using Trimmomatic [10] for quality and adapter trimming with the following parameters: `ILLUMINACLIP:Adapters:2:30:10:8:true`, `HEADCROP:5`, `SLIDINGWINDOW:4:10`, `LEADING:3`, `TRAILING:3`, `MINLEN:30`. Trimmed reads were aligned to the human reference genome (hg19) using BWA with default settings [11]. Pairtools was used to parse and deduplicate the aligned pairs [12], followed by sorting and indexing with samtools [13] and pairix [14]. Final contact matrices were generated using cooler [15] (balanced using the `--balance` option) and stored in `.cool` format for downstream analysis.

Interaction Matrix Preparation and Annotation. Hi-C cool files were converted to sparse interaction matrices at 4-kb resolution. We retained only intra-chromosomal contacts, required a minimum of two interaction counts per pair, and excluded interactions shorter than 4 kb. Sparse matrices from all samples were then merged into a unified matrix. From this merged matrix, individual interaction sites were extracted and defined as anchors. Genomic annotation of anchors was performed using ChIPSeeker, and cancer-associated SNPs were assigned using the most recent GWAS Catalog (EMBL-EBI), filtered for traits containing the term "cancer." Enhancer annotations were obtained from the UCSC EPInteract database (MCF7-specific tracks). Annotated anchors were subsequently recombined with the sparse matrices to annotate each looping pair, thereby assigning genomic features (e.g., promoters, enhancers, cancer SNPs) to each anchor and to each interaction.

Cancer SNP Analysis. We quantified the number of interactions in which either one anchor or both anchors overlapped cancer-associated SNPs. Observed versus expected enrichment was computed by comparing these counts within each interaction class against the background of all interactions. Enrichment values were visualized in GraphPad Prism.

Enhancer Overlap Analysis. For each 4-kb bin, the number of overlapping MCF7-specific enhancers was determined. We then calculated, for each interaction class, the fraction of interactions whose anchors contained a given number of enhancers. The results were displayed as heatmaps generated with ggplot2.

Eperezolid/Radezolid/JQ1 vs DMSO comparison. For each treatment, library-normalized interaction counts were calculated separately for intergenic and intragenic interactions and stratified by interaction type (e.g., promoter–promoter, intron–promoter). Ratios relative to the DMSO control were computed after adding a pseudocount of 0.1 to all values to stabilize very small ratios and avoid division by zero. Log2-transformed treatment/control ratios of all intergenic interaction types with at least 5000 interactions were plotted with GraphPad Prism.

Reactome Pathway Analysis. Reactome pathway enrichment analysis was performed with the ReactomePA R package using gene lists corresponding to promoters/TSS regions engaged in specific loop classes (as shown in the figures). Enrichment significance values (–log10p-values) were plotted in GraphPad Prism.

**AI4Loop application in Single-cell RNA-seq data**

Single-cell RNA-seq (scRNA-seq) drug perturbation datasets were obtained from two publicly available large-scale resources. The first dataset profiled pooled scRNA-seq responses across multiple cancer cell lines (including K562, MCF7, and A549) under diverse small-molecule perturbations [16]. The second dataset provided scRNA-seq drug perturbation profiles for ovarian cancer cell lines (JH0S2, PDC3, and PDC2) across a broad range of compounds [17].

For each dataset, raw single-cell expression matrices were processed according to the original study protocols. Cells were first grouped by cell line and treatment condition. To ensure compatibility with the AI4Loop input format, pseudo-bulk gene counts were converted to FPKM (Fragments Per Kilobase of transcript per Million mapped reads) using gene length annotations from the hg19 reference.

For downstream prediction, AI4Loop was applied directly to the pseudo-bulk FPKM profiles without model re-training. A prediction probability threshold of 0.5 was used to define positive GCIs, providing a stringent criterion to prioritize high-confidence predictions in scRNA-seq, TCGA, and drug perturbation analyses.

### Supplementary Figures

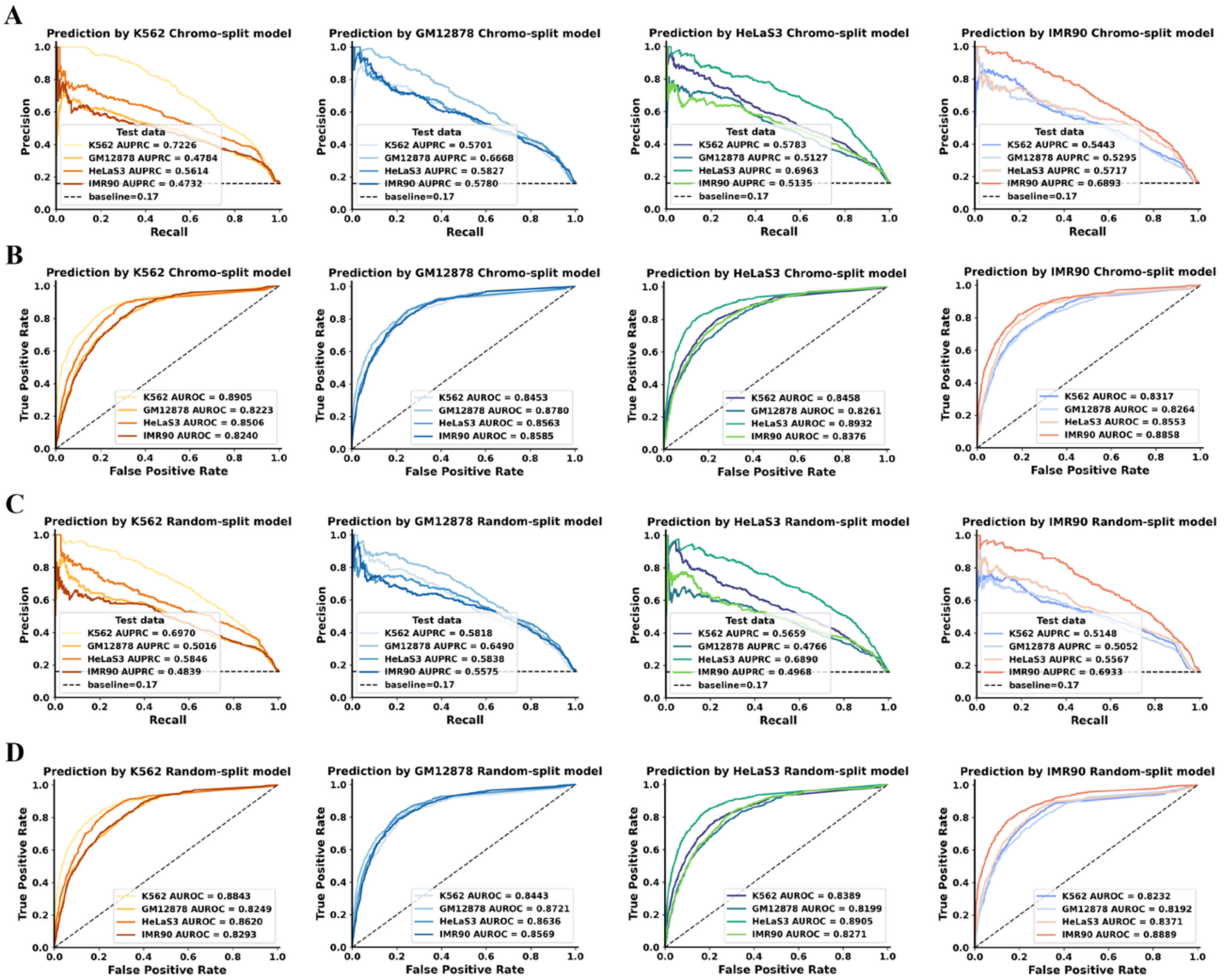


**Fig. S1. Cross-model generalization under chromosome- and random-based splits.** (**A**) **Precision-recall curves obtained using chromosome-split training strategy**. (**B**) Corresponding ROC curves for chromosome-split evaluation. (**C**) Precision-recall curves under random-split training strategy. (**D**) Corresponding ROC curves for random-split evaluation.

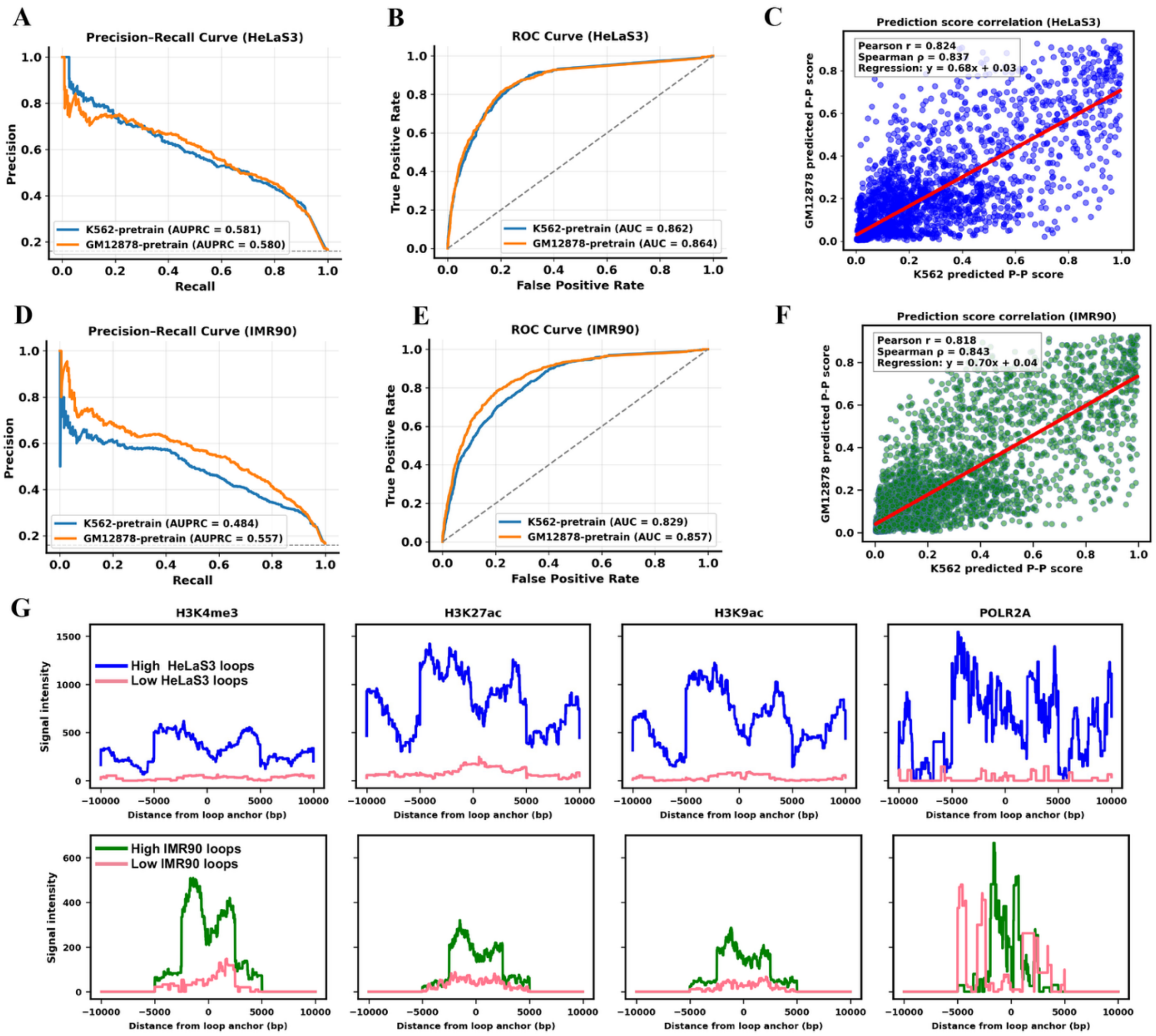


**Fig. S2. Cross-model evaluation and validation of prediction reliability.** (**A-B**) Precision-recall (PR) and ROC curves for HeLaS3 cells using K562-pretrained and GM12878-pretrained models. (**C**) Prediction score correlation in HeLaS3 between K562-pretrained and GM12878-pretrained models. (**D-E**) PR and ROC curves for IMR90 cells using K562-pretrained and GM12878-pretrained models. (**F**) Prediction score correlation in IMR90 between K562-pretrained and GM12878-pretrained models. (**G**) Epigenetic signal profiles of the top 100 (high-prediction) and bottom 100 (low-prediction) predicted loops in the HeLaS2 and IMR90. High-confidence loops were markedly enriched for active histone marks (H3K4me3, H3K27ac, H3K9ac) and transcriptional regulators (POLR2A).

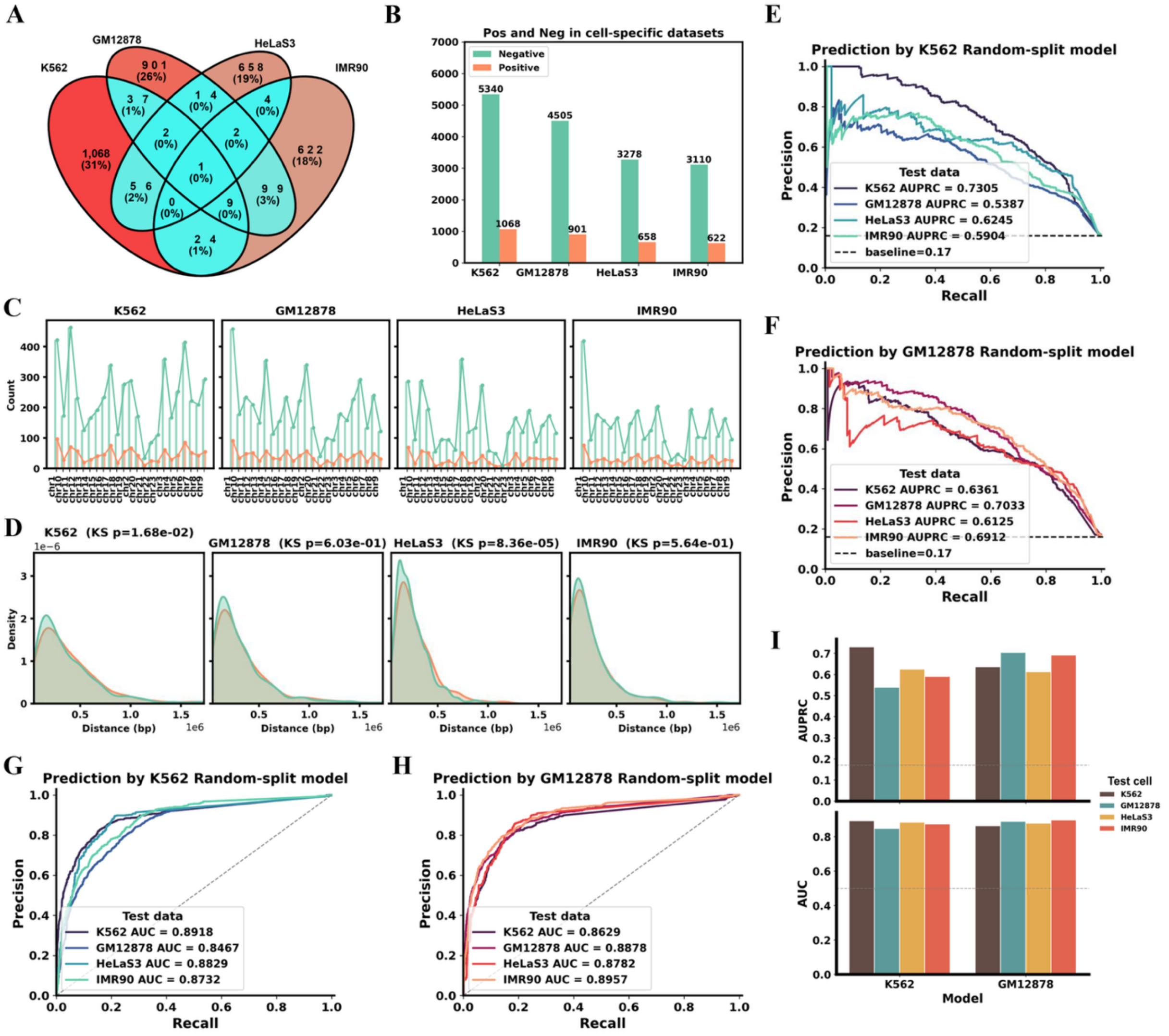


**Fig. S3. AI4Loop detects cell-type-specific gene-centered chromatin interactions (GCIs) across multiple cell types.** (**A**) Venn diagram showing the number of GCIs unique to each cell type (K562, GM12878, HeLaS3, and IMR90) based on a 10 kb anchor-matching criterion. (**B**) Bar plot summarizing the number of positive (cell-type-specific loops) and distance-matched negative samples used to construct cell-specific benchmarking datasets. (**C**) Chromosome-wise distribution of positive and negative samples for cell-type-specific datasets across the four cell types. (**D**) Genomic distance density plots demonstrating successful distance matching between positive and negative samples for each cell type (Kolmogorov–Smirnov test p-values shown). (**E-F**) Precision–recall curves showing cross-cell prediction performance when using the K562-trained model (E) and GM12878-trained model (F) to predict cell-type-specific loops from all four cell types. (**G-H**) Corresponding ROC curves demonstrating cross-cell generalization using the K562-trained (G) and GM12878-trained (H) models on cell-type-specific datasets. (**I**) Summary bar charts comparing AUPRC and AUC across models and test cell types.

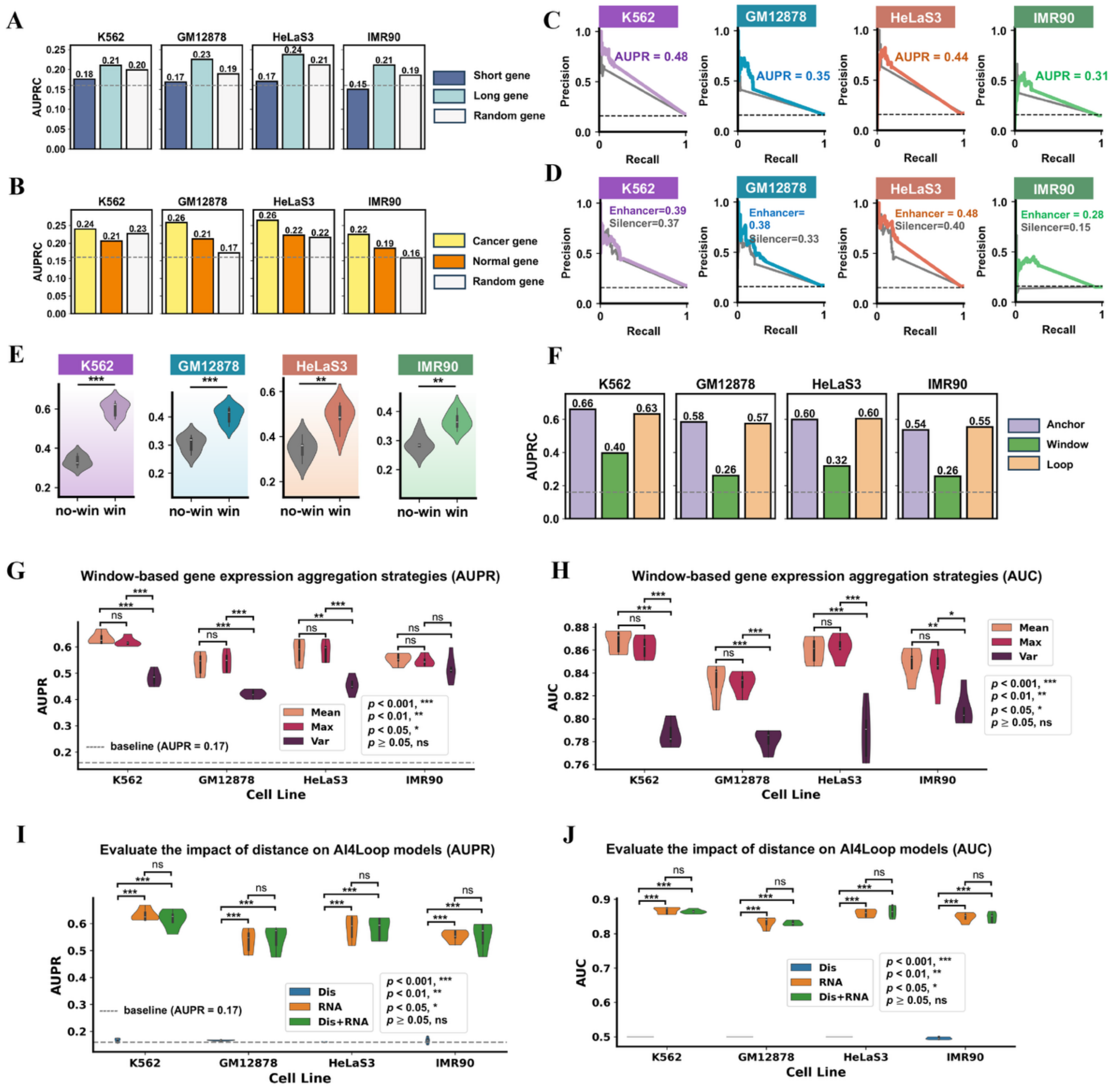


**Fig. S4. Feature design and attribution analysis of the AI4Loop framework.** (**A**) Comparison of models trained by gene expression feature sets from 1,000 short genes and 1,000 long genes. Randomly select the same number of genes as the control group. (**B**) Comparison of models trained by gene expression feature sets from 1,108 cancer genes and 1,108 normal genes. Randomly select the same number of genes as the control group. (**C**) Comparison of models trained by high-level gene expression of 1,000 genes and gene expression feature sets of randomly selected 1,000 genes (gray). (**D**) Comparison of models trained by 2,000 enhancer genes and 2,000 silencer genes (gray). The enhancer genes were defined as genes colocalized with H3K27ac. The silencer genes were defined as genes colocalized with H3K9me3. (**E**) Comparison of models trained by no-window-based (no-win) and window-based (win) methods. (*$p < 0.01$, **$p < 0.001$). (**F**) Comparison of models trained by gene expression feature sets from anchors, window and loop. The window means the region between two anchors. The loop was defined as the region of two anchors and window. (**G-H**) Comparison of window-based gene expression aggregation strategies using mean, maximum, and variance, evaluated by AUPR (G) and AUC (H) across cell lines. **(I-J)** AUPR and AUC comparison of AI4Loop models trained using genomic distance alone (Dis), RNA-seq features alone (RNA), or their combination (Dis+RNA) across four cell lines.

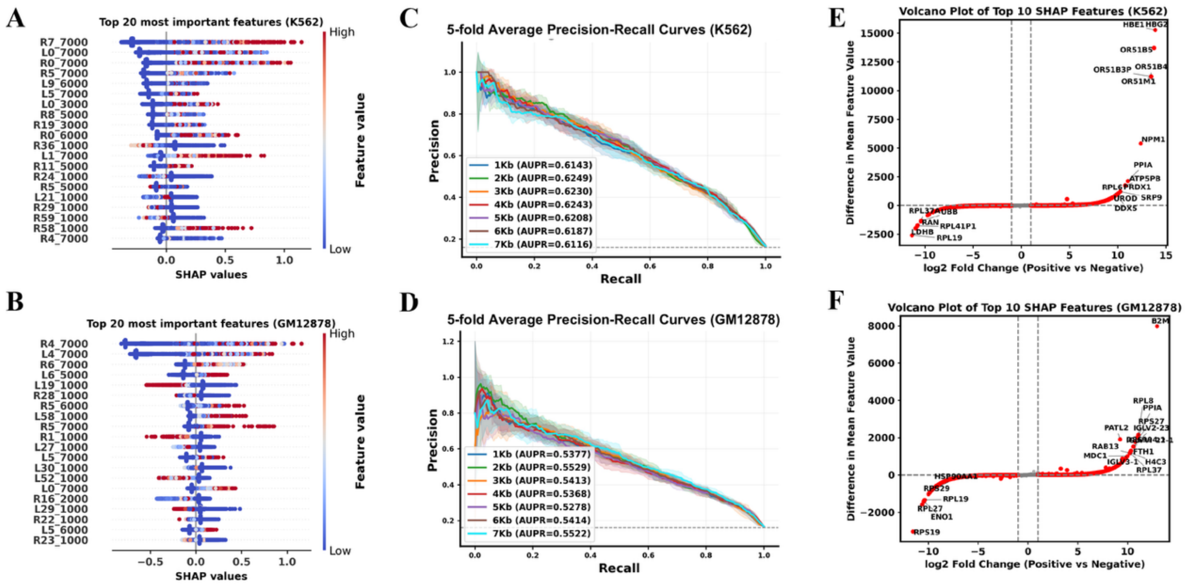


**Fig. S5. Feature attribution analysis of the AI4Loop framework.** (**A-B**) SHAP summary plots of the top 20 most important features for K562 (A) and GM12878 (B). Features are ranked by mean absolute SHAP values, with color indicating feature magnitude (low to high). (**C-D**) Five-fold cross-validated precision-recall curves for different anchor window sizes (1-7 kb) in K562 (C) and GM12878 (D). (**E-F**) Volcano plots of the top 20 SHAP-ranked features for K562 (E) and GM12878 (F), showing $\log_2$ fold change between positive and negative samples versus differences in mean feature values.

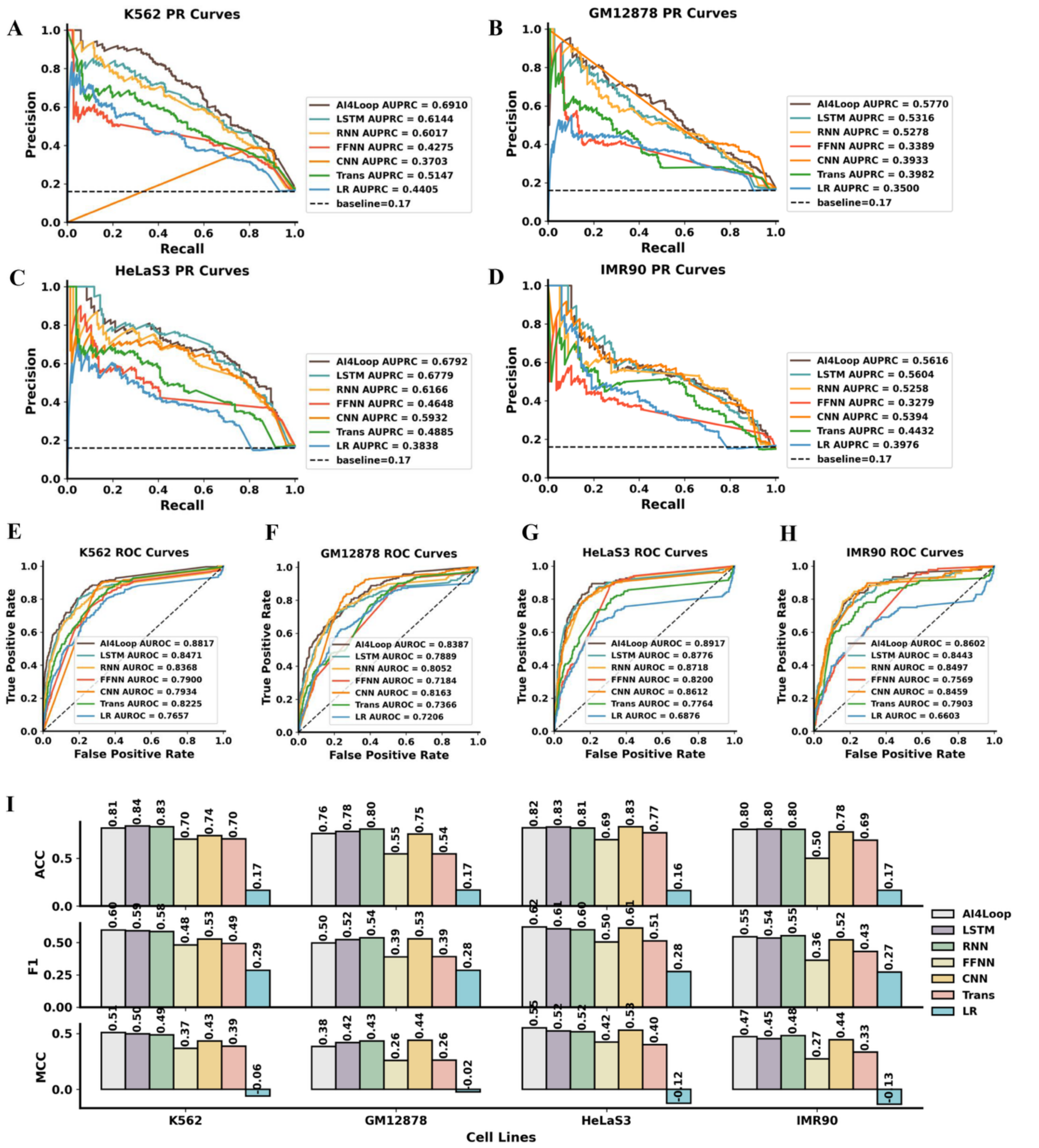


**Fig. S6. Performance comparison of AI4Loop with alternative machine learning models across four cell lines.** (**A-D**) Precision-Recall (PR) curves and corresponding AUPRC values. (**E-H**) Receiver Operating Characteristic (ROC) curves and corresponding AUROC values. (**I**) Bar plots showing accuracy (ACC), F1-score (F1), and Matthews correlation coefficient (MCC). Evaluated algorithms include AI4Loop, Transformer (Trans), Convolutional Neural Network (CNN), Feedforward Neural Network (FFNN), Recurrent Neural Network (RNN), Long Short-Term Memory (LSTM), and Logistic Regression (LR). The baseline for PR curves corresponds to the proportion of positive samples in each dataset.

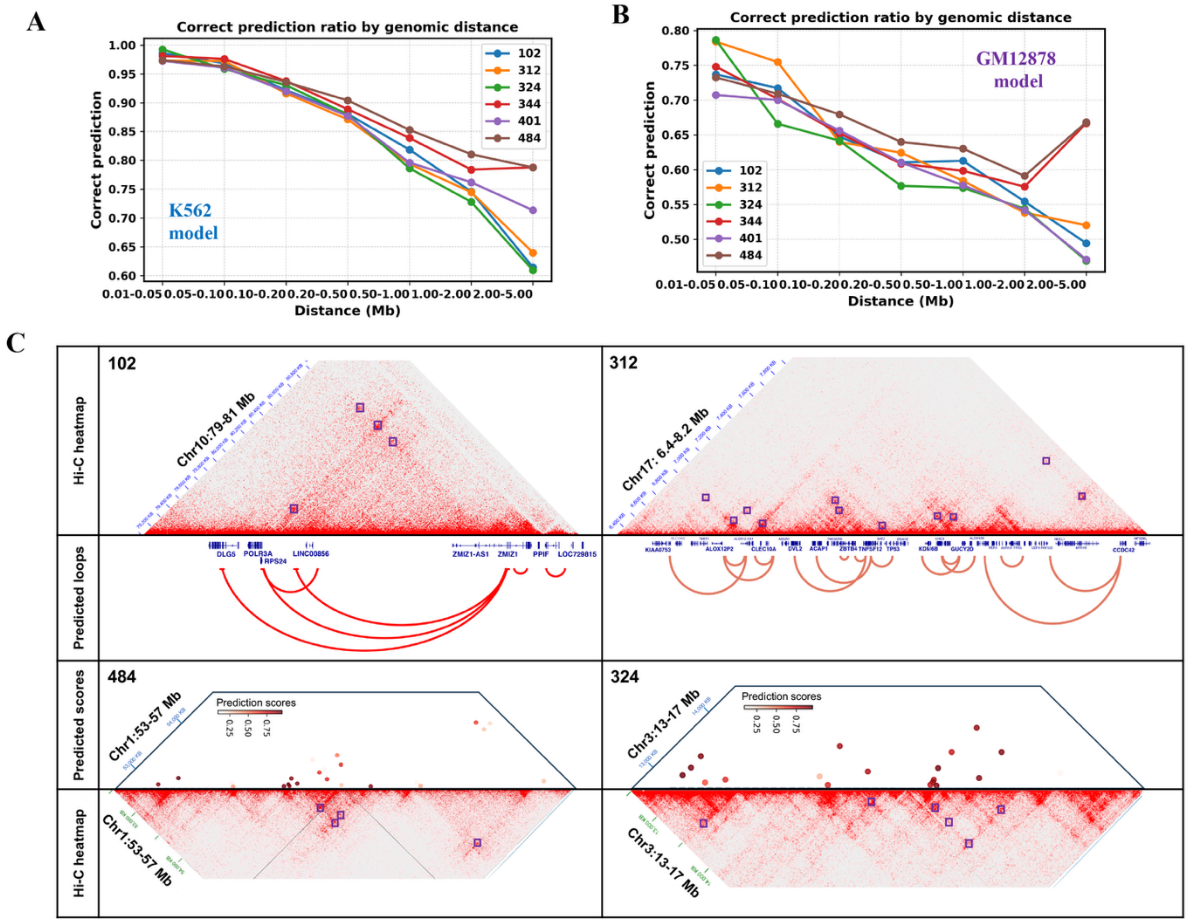


**Fig. S7. Distance-stratified evaluation of AI4Loop prediction accuracy in CLL samples.** (**A-B**) Correct prediction ratios stratified by genomic distance for CLL samples using AI4Loop models trained on K562 (A) and GM12878 (B). Predictions were grouped into distance bins (0.01-5 Mb). (**C**) Representative genomic regions illustrating concordance between high-confidence AI4Loop predictions and experimentally observed Hi-C interaction patterns in CLL samples. Upper panels show Hi-C contact heatmaps vs. predicted chromatin loops, while lower panels display predicted scores across selected loci vs. Hi-C contact heatmaps.

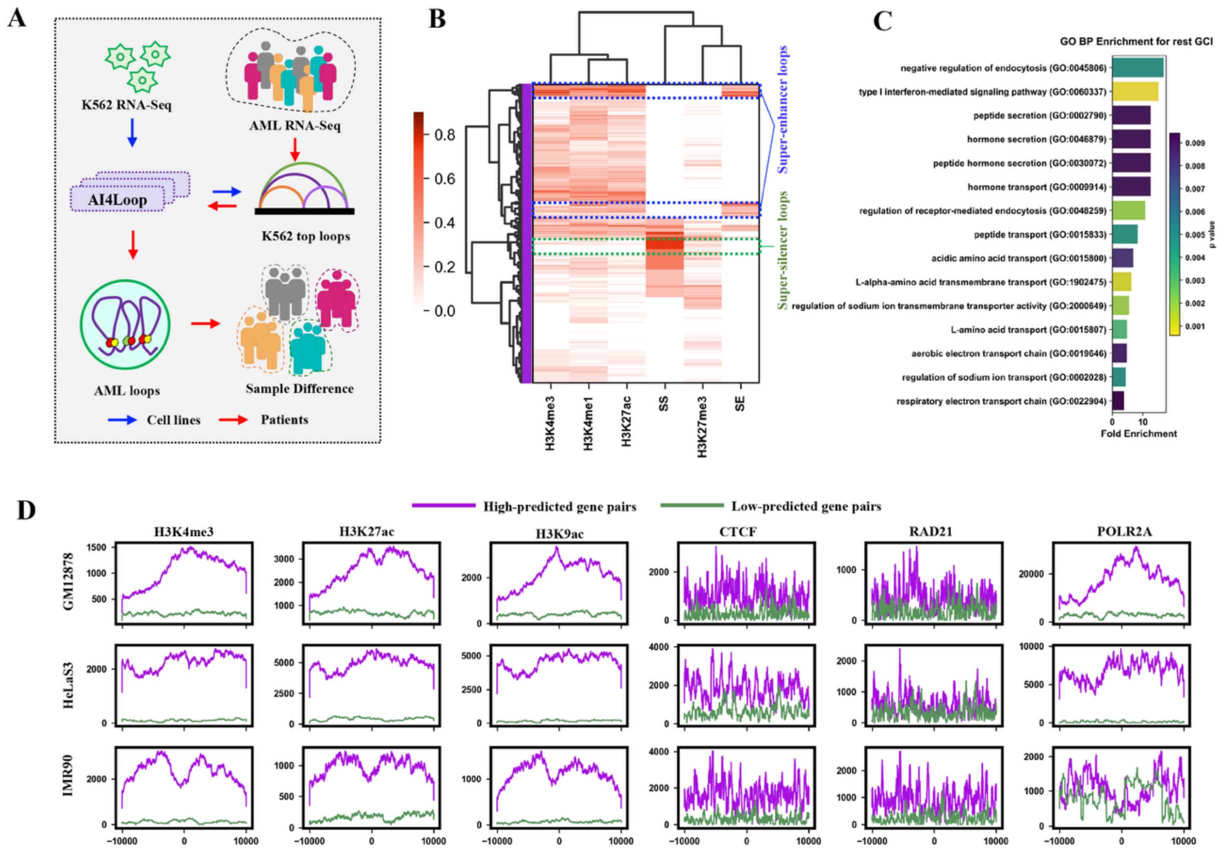


**Fig. S8. AI4Loop model was used to reveal the chromatin interaction differences between different Acute Myeloid Leukemia (AML) samples and subtypes.** (**A**) Schematic diagram showing AI4Loop model mining AML specific GCIs. (**B**) Hierarchical clustering high predicted loops of K562 based on the enrichment of epigenetic modification signals. SS: super-silencer, SE: super-enhancer. (**C**) Gene Ontology (GO) functional analysis of the rest GCIs (not AML GCIs) (**D**) The distribution of epigenetic modification signals at loop anchors with high and low prediction probabilities for different cell lines.

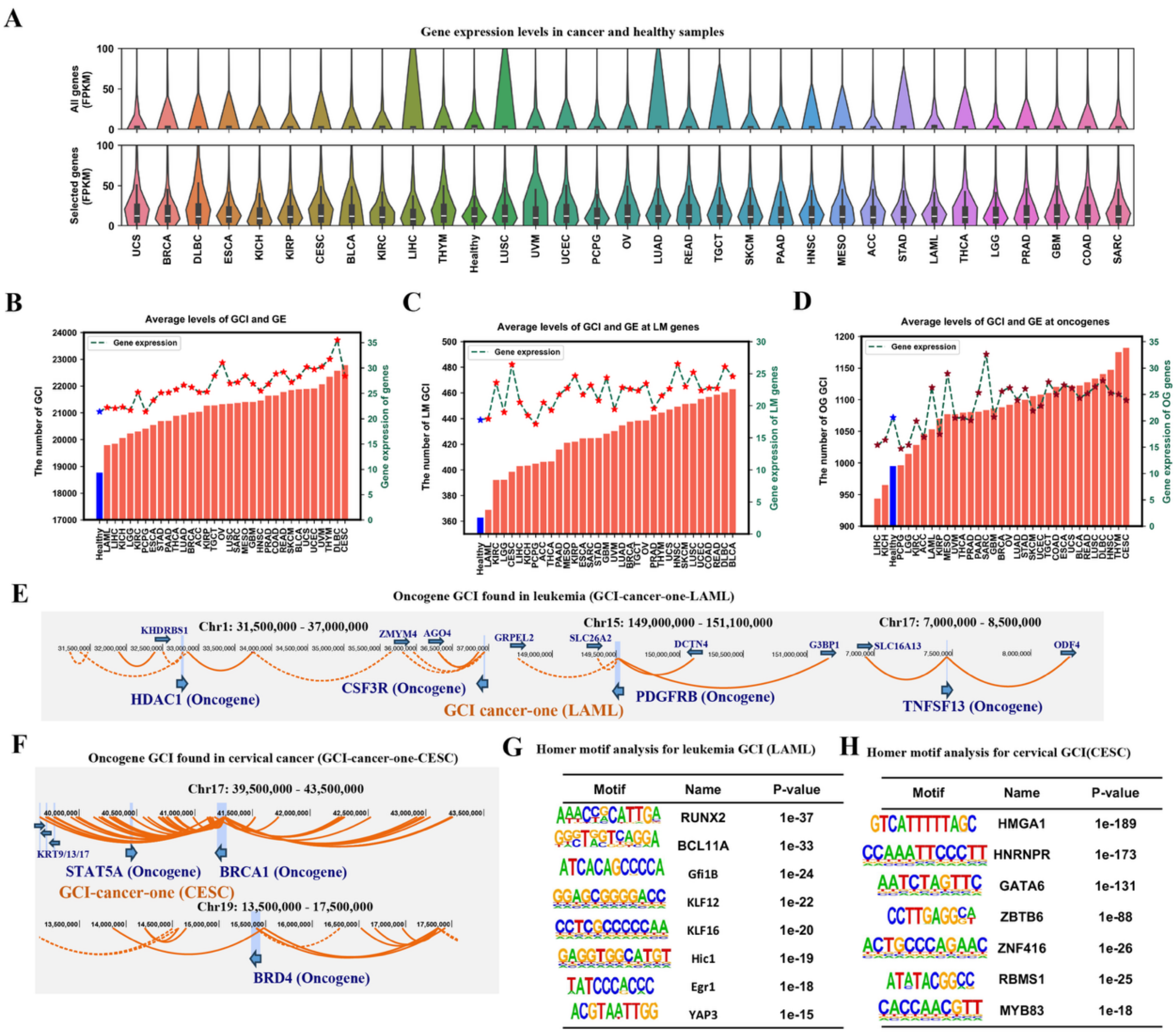


**Fig. S9**. **AI4loop reveals that cancers have increased gene-centered chromatin interactions (GCIs) as compared with healthy samples.** (**A**) The gene expression level (FPKM) of 55,420 genes and 5,338 selected genes. (**B-D**) The number of predicted GCIs and average gene expression levels in cancer and healthy samples for all gene pairs (B), landmark gene pairs (C), and oncogene pairs (D). (**E-F**) GCI-cancer-one (LAML and CESC) at oncogenes. (**G-H**) Enriched motifs of GCI-cancer-one (LAML and CESC).

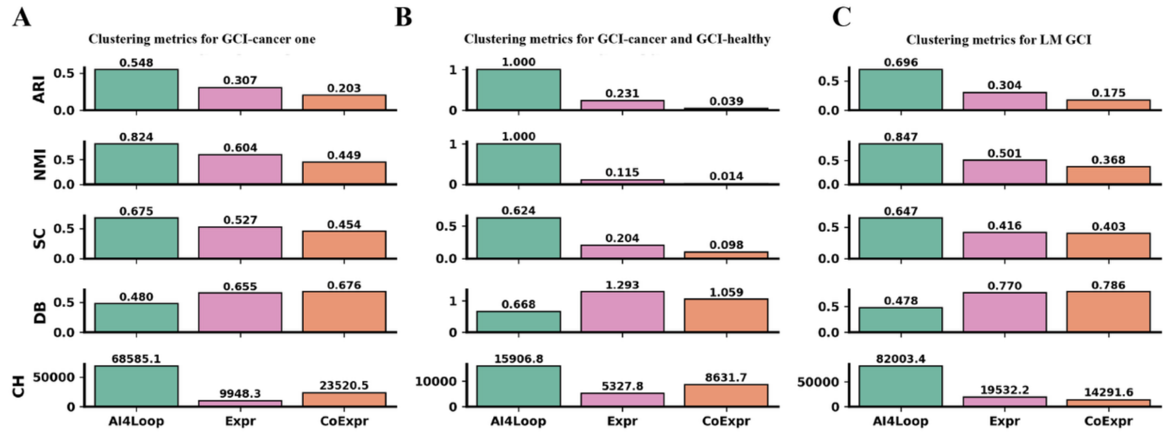


**Fig. S10. Quantitative evaluation of clustering performance for cancer and healthy sample separation using gene-centered chromatin interaction features.** (**A**) Clustering metrics for samples represented by GCI-cancer-one features. (**B**) Clustering metrics for samples represented by GCI-cancer and GCI-healthy features. (**C**) Clustering metrics for samples represented by landmark gene (LM) GCI features. For each feature set, clustering performance was compared across AI4Loop-predicted GCI probabilities (AI4Loop), gene expression (Expr), and gene co-expression (CoExpr) using the adjusted Rand index (ARI), normalized mutual information (NMI), silhouette coefficient (SC), Davies–Bouldin index (DB), and Calinski–Harabasz score (CH). Higher ARI, NMI, SC, and CH values indicate better clustering performance, whereas lower DB values indicate better clustering compactness and separation.

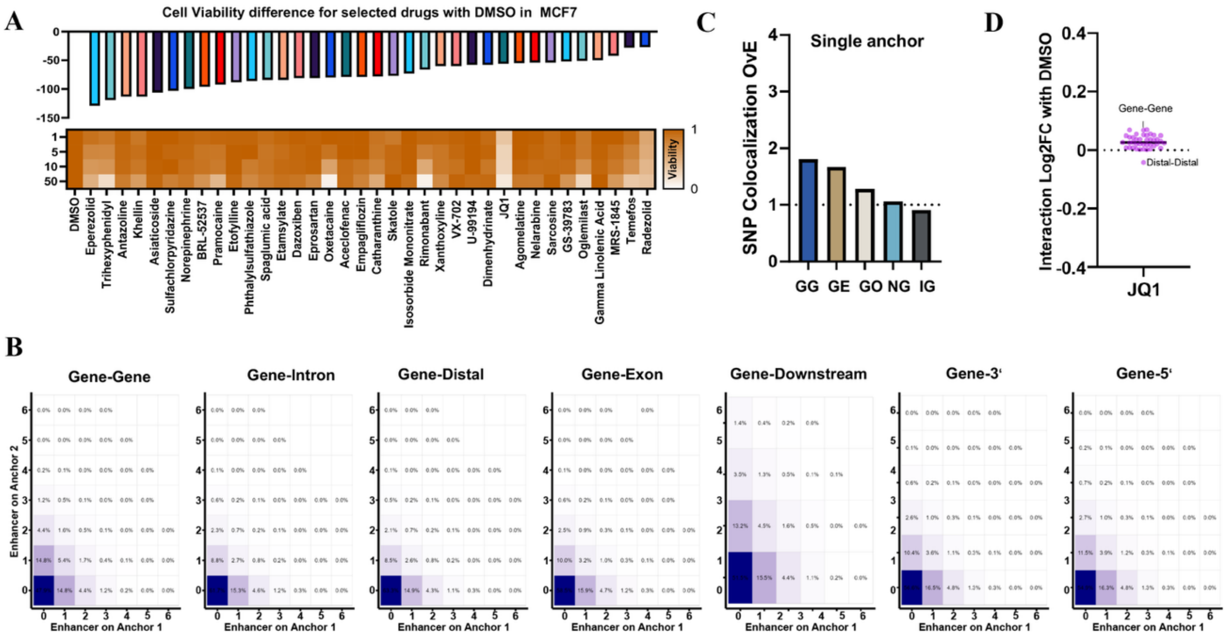

**Fig. S11. Drug-induced perturbation of gene-centered chromatin interaction (GCI) and functional annotation in MCF7 cells.** (**A**) Cell viability effects (heatmap) and predicted GCIs (bar chart) for selected compounds predicted to disrupt GCIs. (**B**) Proportion of gene-related intergenic interactions containing a defined number of MCF7-specific enhancers at each anchor. (**C**) Observed versus expected enrichment of cancer-associated SNPs overlapping at least one anchor in gene-gene (GG), gene-enhancer (GE), gene-other (GO; anchors lacking gene or enhancer annotation), non-gene (NG), and intragenic (IG) interactions. (**D**) Distribution of library-normalized $\log_2$ fold change (Log2FC) in interaction strength under JQ1 perturbation relative to DMSO for different loop categories, comparing gene-gene and distal-distal interactions.

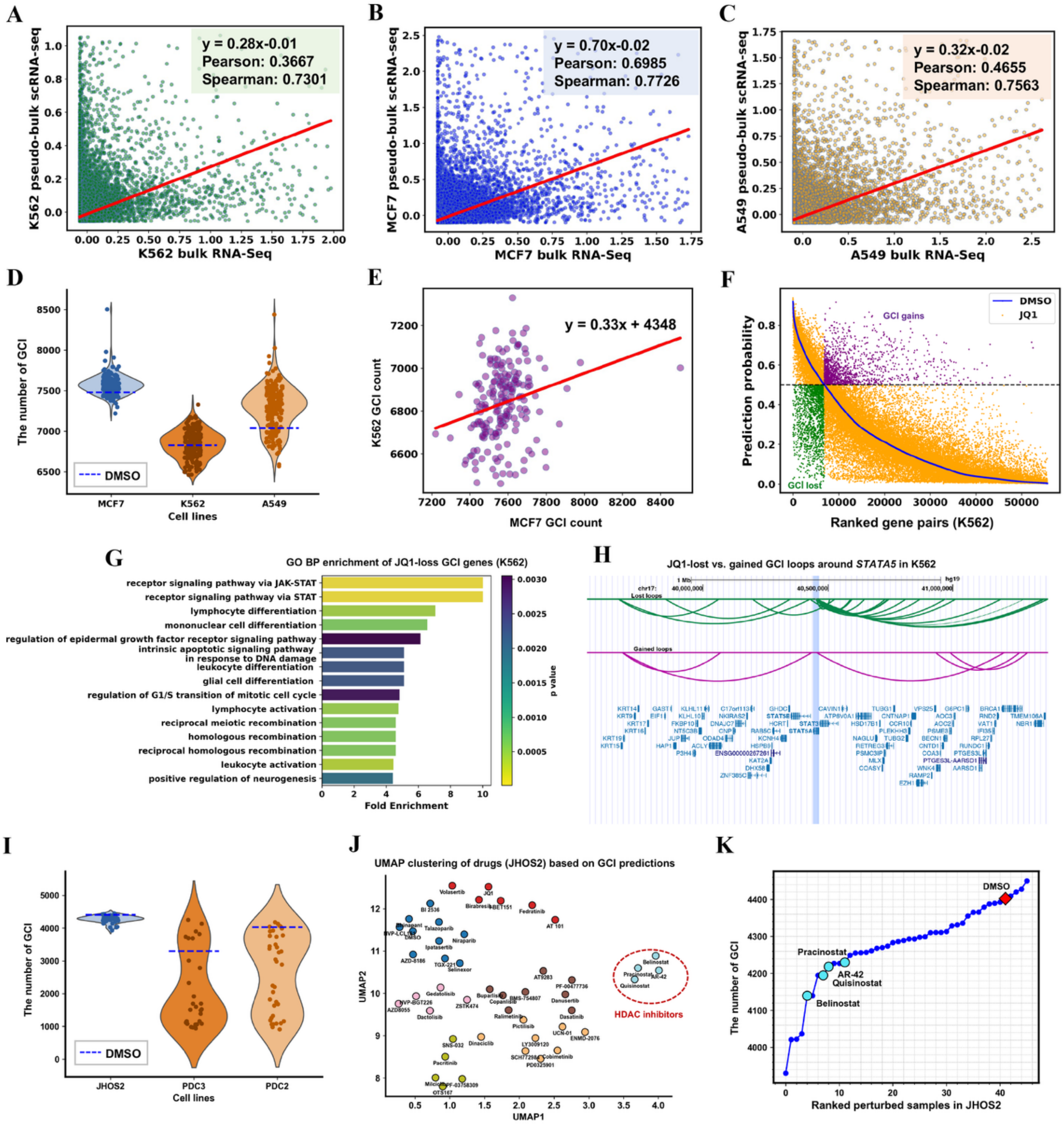


**Fig. S12. Application of AI4Loop to single-cell RNA-seq drug perturbation datasets.** (**A-C**) Correlation between pseudo-bulk scRNA-seq expression profiles and matched bulk RNA-seq for K562, MCF7, and A549 cells. (**D**) Predicted numbers of GCIs in MCF7, K562, and A549 under DMSO control versus drug-treated conditions using the pre-trained GM12878 model. (**E**) Positive correlation between predicted GCI counts in K562 and MCF7 across perturbations. (**F**) Ranked prediction probabilities for K562 under JQ1 treatment relative to DMSO, revealing clear separation of GCI gains and losses. (**G**) GO Biological Process enrichment analysis of JQ1-induced GCI losses in K562, highlighting pathways related to JAK-STAT signaling, lymphocyte differentiation, and regulatory programs consistent with known JQ1 biology. (**H**) Genome browser-style visualization of lost and gained GCI loops around the STAT gene locus in K562 upon JQ1 treatment. (**I**) Predicted numbers of GCI in JHOS2, PDC3, and PDC2 cells under DMSO and drug perturbation using scRNA-seq profiles. (**J**) UMAP clustering of drug perturbations in JHOS2 based on AI4Loop-predicted GCI profiles, revealing coherent grouping of HDAC inhibitors. (**K**) Ranking of perturbed JHOS2 samples based on total predicted GCI counts, with HDAC inhibitors associated with pronounced GCI losses.

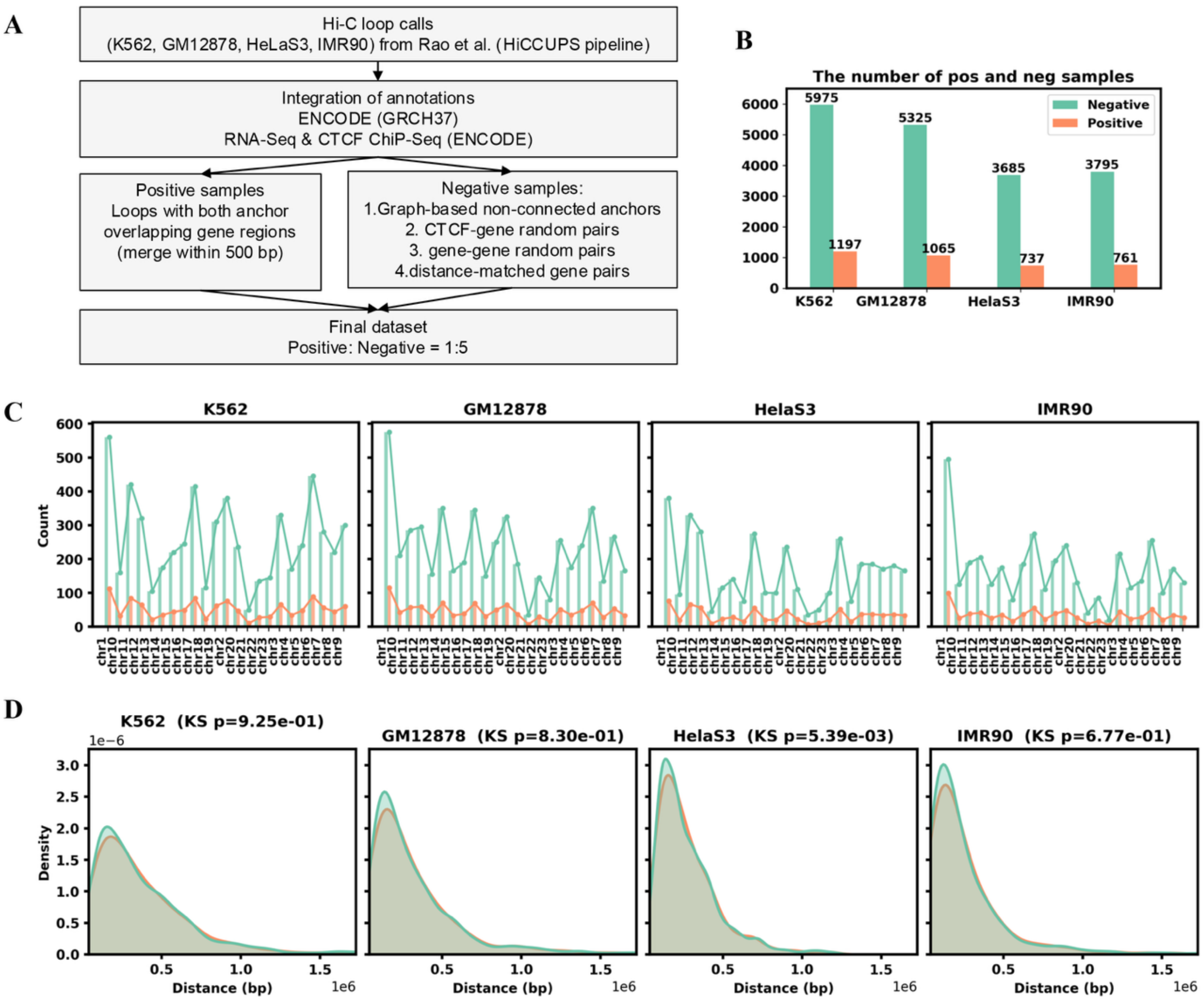


**Fig. S13. The datasets construction across four cell lines.** (**A**) Definition of positive and negative samples. (**B**) The number of positive and negative samples across four datasets. (**C**) The number of positive and negative samples in each chromosome across four datasets. (**D**) The distance distribution of positive and negative set in four datasets of K562, GM12878, HeLaS3, IMR90 cell lines. KS: Kolmogorov-Smirnov test.
